# Targeted sequencing of *Enterobacterales* bacteria using CRISPR-Cas9 enrichment and Oxford Nanopore Technologies

**DOI:** 10.1101/2024.06.26.600727

**Authors:** Hugh Cottingham, Louise M. Judd, Jessica A. Wisniewski, Ryan R. Wick, Thomas D. Stanton, Ben Vezina, Nenad Macesic, Anton Y. Peleg, Iruka N. Okeke, Kathryn E. Holt, Jane Hawkey

## Abstract

Sequencing DNA directly from patient samples enables faster pathogen characterisation compared to traditional culture-based approaches, but often yields insufficient sequence data for effective downstream analysis. CRISPR-Cas9 enrichment is designed to improve yield of low abundance sequences but has not been thoroughly explored with Oxford Nanopore Technologies (ONT) for use in clinical bacterial epidemiology. We designed CRISPR-Cas9 guide RNAs to enrich for the human pathogen *Klebsiella pneumoniae*, by targeting multi-locus sequence type (MLST) and transfer RNA (tRNA) genes, as well as common antimicrobial resistance (AMR) genes and the resistance-associated integron gene *intI1*. We validated enrichment performance in bacterial isolates before comparing enriched and unenriched sequencing of three human faecal samples spiked with *K. pneumoniae* at varying abundance. Enriched sequencing generated 56x and 11.3x the number of AMR and MLST reads respectively compared to unenriched sequencing and required approximately one third of the computational storage space. Targeting the *intI1* gene often led to detection of 10-20 proximal resistance genes due to the long reads produced by ONT sequencing. We demonstrated that CRISPR-Cas9 enrichment combined with ONT sequencing enabled improved genomic characterisation outcomes over unenriched sequencing of patient samples. This method could be used to inform infection control strategies by identifying patients colonised with high-risk strains.

## Introduction

Effective and rapid characterisation of AMR bacterial pathogens is crucial for improving patient outcomes and containing outbreaks in hospital settings. Current gold-standard characterisation methods, such as whole genome sequencing, MALDI-TOF mass spectrometry and antimicrobial susceptibility testing, rely on time-consuming bacterial culture (Lagier et al. 2015). Modern high-throughput sequencing technologies, such as Illumina and Oxford Nanopore Technologies (ONT), have the capacity to vastly improve characterisation speed by bypassing bacterial culture and sequencing pathogenic DNA directly from patient samples. However, many sample types, including faecal, saliva, nasal and vaginal specimens, contain pathogen DNA at <10% abundance of total DNA (Huttenhower et al. 2012; Marotz et al. 2018; Yang et al. 2020). This often leads to insufficient pathogen sequence data for effective characterisation. Several methods have been developed to enrich for low abundance pathogen DNA prior to sequencing, but all have significant limitations. Host DNA depletion methods (e.g. saponin enrichment and CpG methylated DNA removal) require fresh, unfrozen samples to be effective and are not selective for specific bacteria (Hasan et al. 2016; Thoendel et al. 2016; Charalampous et al. 2019; Street et al. 2020). Amplicon sequencing methods such as selective whole genome amplification can take days to complete and often cannot target multiple different pathogens at once (Clarke et al. 2017; Cocking et al. 2020). Hybrid capture-based methods suffer from high financial costs and a lack of target flexibility (Hodges et al. 2007; Brown et al. 2015). ONT’s adaptive sampling shows promise for depleting unwanted DNA during sequencing, but has thus far been unable to substantially increase absolute numbers of target sequences and often leads to premature flowcell degradation (Loose et al. 2016; Edwards et al. 2019; Kovaka et al. 2021; Payne et al. 2021; Ulrich et al. 2024).

CRISPR-Cas9 enrichment allows the selective sequencing of DNA fragments containing a chosen 23 bp target sequence. This approach is applied immediately following DNA extraction and begins with the removal of terminal phosphate groups from genomic DNA, preventing phosphate-dependent sequencing adapters from ligating to DNA molecules so they will not be available for sequencing (**Figure 1**). A pool of CRISPR-Cas9 guide RNAs (guides) is then used to direct Cas9 cleavage of sequences with a complementary sequence, exposing internal phosphate groups so that sequencing adapters can then selectively ligate to cleaved molecules, making them available for sequencing (**Figure 1**).

**Figure 1.**
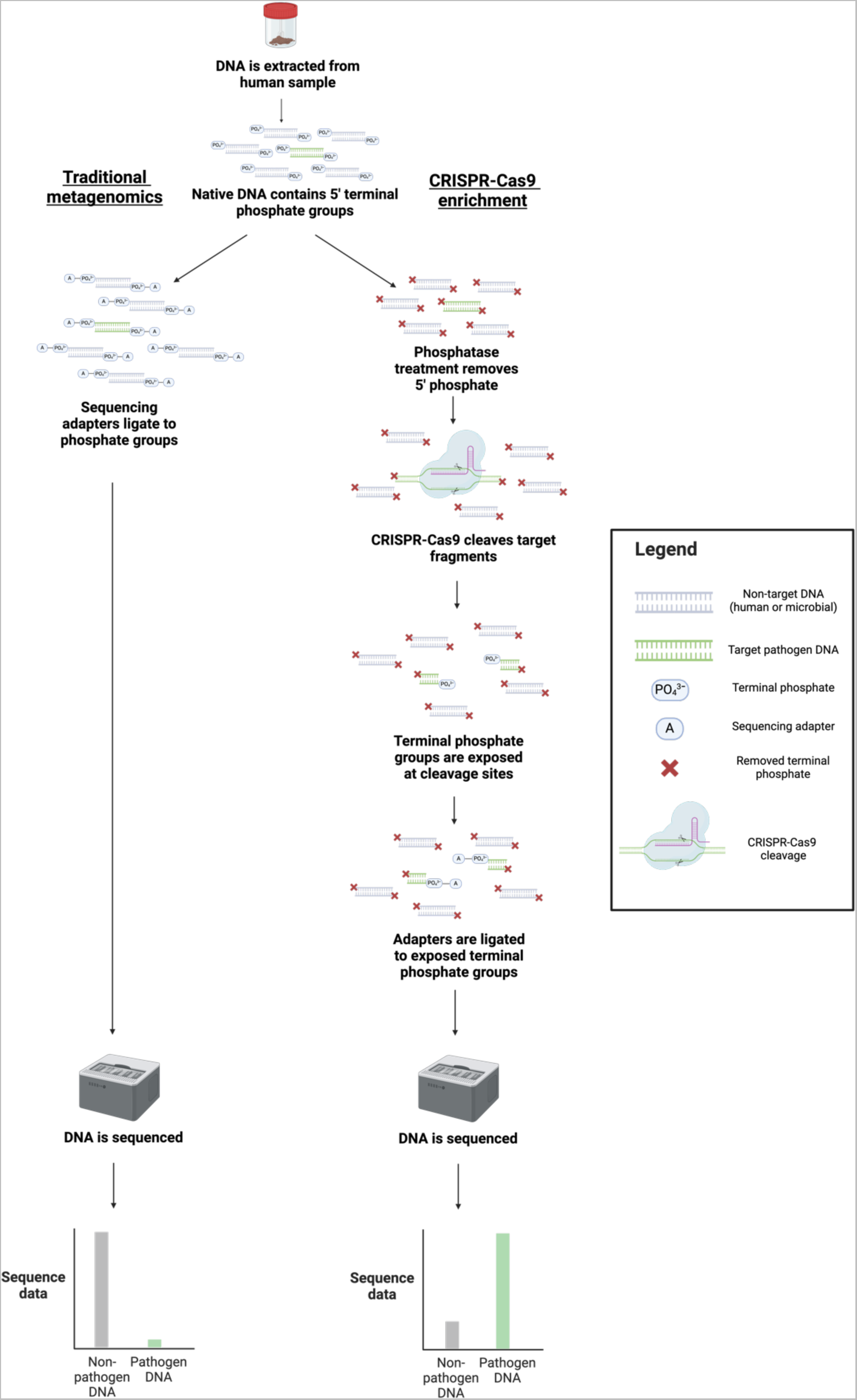
Library preparation differences between unenriched and CRISPR-Cas9 enriched sequencing. During unenriched sequencing, sequencing adapters are ligated to native terminal phosphate groups on DNA molecules to allow for sequencing. During CRISPR-Cas9 enrichment, native phosphate groups are removed from all DNA molecules so that adapters cannot ligate. CRISPR-Cas9 is then used to cleave molecules of interest, exposing their terminal phosphate groups and allowing for specific adapter ligation and sequencing.

The first published use of CRISPR-Cas9 for enrichment of bacterial DNA was employed by Quan et al, who targeted 127 AMR genes in *Staphylococcus aureus* and *Enterococcus faecium* (Quan et al. 2019). While this study demonstrated effective detection of low abundance pathogens, it also highlighted the limitations of using short read sequencing. Illumina short read sequencing requires adapters be present at both ends of the molecule, therefore requiring a minimum of two guide RNAs to cleave at nearby sites to create a molecule with adapters at both ends. Short read lengths also generate minimal information on the genetic context of the target genes (Quan et al. 2019). ONT platforms allow for theoretically unlimited read lengths, which could vastly improve the amount of genetic information obtained from each enrichment site. Previous bacterial studies combining CRISPR-Cas9 enrichment with long read sequencing showcased these benefits, but primarily focused on AMR genes (Baltrus et al. 2019; Sajuthi et al. 2020; Serpa et al. 2022).

Here, we chose to focus on enrichment of *K. pneumoniae* and closely related species comprising the *K. pneumoniae* species complex (KpSC) (Mariappan et al. 2017; Wyres et al. 2020), which are associated with high levels of AMR and virulence that can lead to severe cases of sepsis, pneumonia and urinary tract infections (Paczosa and Mecsas 2016). Additionally, this species belongs to the *Enterobacterales,* which accounts for a large proportion of carbapenem-resistant infections in hospitals (Aslan and Paterson 2024). We present an implementation of CRISPR-Cas9 enrichment and ONT sequencing targeting transfer RNA (tRNA), AMR and multi-locus sequence type (MLST) genes in *K. pneumoniae* and other *Enterobacterales* pathogens. We demonstrate successful enrichment of target sequences using DNA extracted from (i) bacterial isolates, (ii) artificial isolate mixtures, and (iii) spiked human faecal samples. In addition, we provide an effective computational workflow for obtaining key characterisation outcomes, including sequence type (ST) and AMR allelic variants, after sequencing.

## Results

### CRISPR-Cas9 guides showed high conservation for target genes and species

To facilitate enrichment of *Enterobacterales* pathogens, we selected 18 tRNA gene sequences from *K. pneumoniae* strain SGH10 as enrichment targets. We chose tRNA genes due to their distribution around the chromosome and conservation within KpSC and *Enterobacterales.* Our CRISPR-Cas9 guides were designed using a pairing approach, comprised of two guides per gene conserved on opposing strands (**Supp. Table 1**, see **Methods**). To predict how our guides would perform in *Enterobacterales,* we aligned guides to all genomes classified as *Enterobacterales* in a dereplicated version of the Genome Taxonomy Database (GTDB) (version R95, n=11,339 genomes, see **Methods**) (Parks et al. 2018). For each guide, we summarised the proportion of genomes per genus harbouring an exact sequence match, which in principle enables targeted enrichment from the corresponding tRNA. One third of genera (35.6%, 89/250) had at least 10 sites with a matching guide sequence in ≥75% of genomes for that genus, while 6.4% (16/250) had at least 30 such sites (**Figure 2, Supp. Table 2**). Most of these 16 genera were in the *Enterobacteriaceae* family, which contains multiple genera of clinical interest such as *Escherichia, Citrobacter, Enterobacter* and *Klebsiella* (**Figure 2B**). In *Klebsiella*, 34/36 guides were highly conserved in 8/11 species, including *K. pneumoniae* (**Supp.** Figure 1).

**Figure 2.**
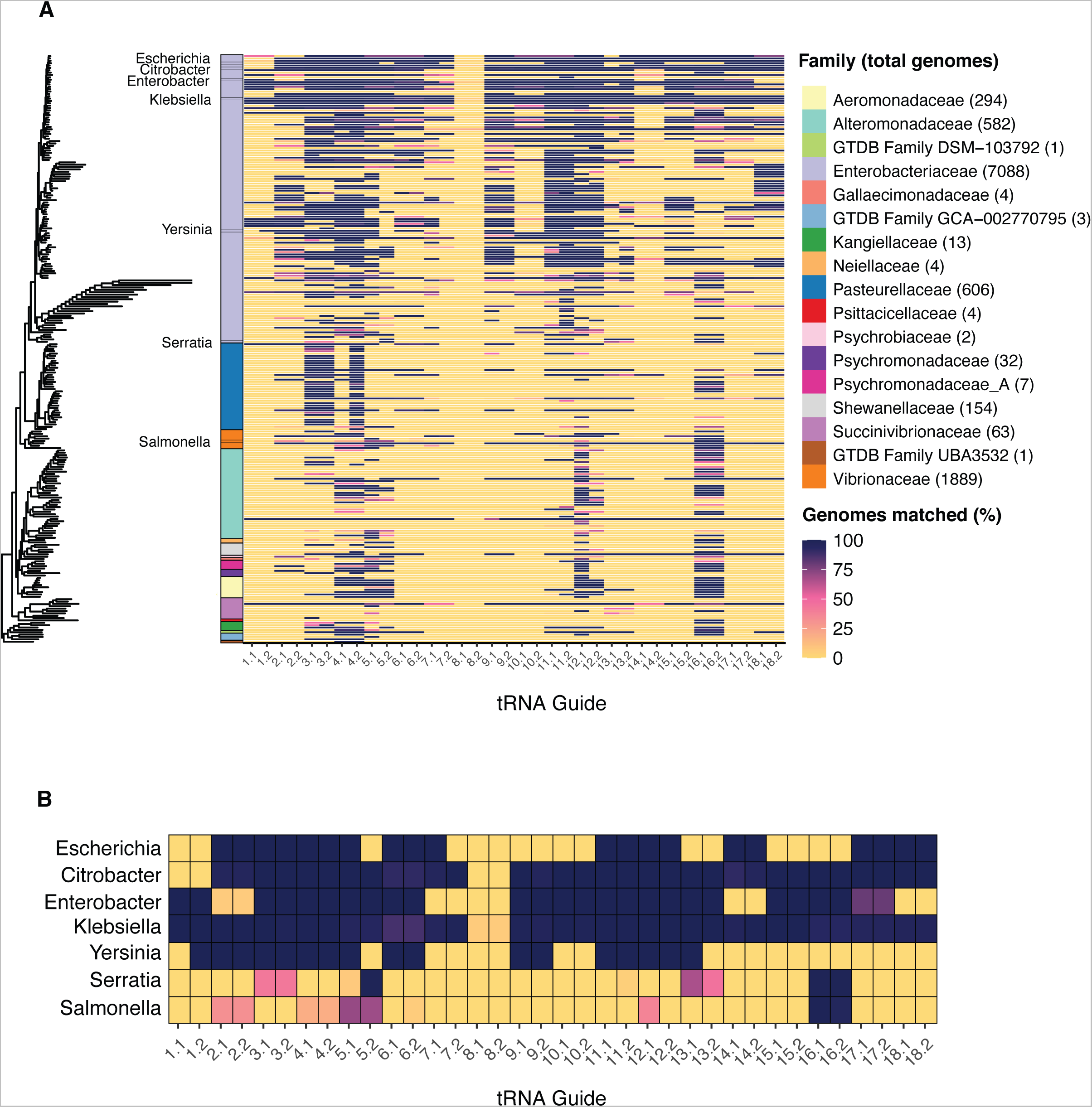
Conservation of tRNA guides across *Enterobacterales*. **A)** Neighbour-joining tree of representative genomes from all genera in GTDB R95 classified as *Enterobacterales* (one genome per genus, n=250 genera) (Parks et al. 2018; Lee S. Katz 2019). The colour spectrum of the heatmap shows the proportion of genomes matched to guide sequences in a dereplicated version of the full GTDB database for each genus (n=11,339 total *Enterobacterales* genomes, genome count for each family shown in brackets). The colour bar to the left of heatmap shows the GTDB-defined family of each genus. **B)** Guide conservation in notable *Enterobacterales* pathogens.

**Table 1.** Sequencing and enrichment statistics following CRISPR-Cas9 enriched and unenriched sequencing of human faecal samples spiked with varying abundances of *K. pneumoniae* strain INF298 (Genbank: GCA_904864465.1).

| Sample | *K. pneumoniae*<br>spike in<br>(CFU/g) | *K. pneumoniae*<br>relative<br>abundance<br>(qPCR) (%) | Enrichment | Library<br>input<br>(ng) | Sequencing<br>duration (hrs) | *K. pneumoniae*-<br>classified bp (per ng<br>DNA input) | *K. pneumoniae* strain<br>INF298 genome coverage<br>(% bases > 0x depth) | *K. pneumoniae*<br>MLST reads | *bla*<sub>CTX-M-15</sub><br>reads | *bla*<sub>OXA-48</sub><br>reads | Class 1<br>integron<br>coverage<br>(% >= 1x<br>depth) |
| --- | --- | --- | --- | --- | --- | --- | --- | --- | --- | --- | --- |
| 1 | 0 | 0.11 | enriched | 84 | 10 | 161 | 0.17 | 0 | 0 | 0 | 0.0 |
| 1 | 0 | 0.11 | unenriched | 20 | 10 | 100 | 0.01 | 0 | 0 | 0 | 0.0 |
| 1 | 4x10<sup>6</sup> | 0.31 | enriched | 78 | 10 | 1,669 | 1.97 | 1 | 1 | 0 | 4.5 |
| 1 | 4x10<sup>6</sup> | 0.31 | unenriched | 12 | 10 | 1,231 | 0.2 | 0 | 0 | 0 | 0.0 |
| 1 | 4x10<sup>7</sup> | 2.14 | enriched | 90 | 40 | 7,280 | 5.63 | 10 | 3 | 5 | 13.9 |
| 1 | 4x10<sup>7</sup> | 2.14 | unenriched | 108 | 40 | 14,191 | 16.16 | 3 | 0 | 0 | 0.0 |
| 2 | 0 | 0.01 | enriched | 48 | 10 | 66 | 0 | 0 | 0 | 0 | 0.0 |
| 2 | 0 | 0.01 | unenriched | 66 | 10 | 40 | 0 | 0 | 0 | 0 | 0.0 |
| 2 | 4x10<sup>6</sup> | 0.27 | enriched | 132 | 10 | 1,115 | 2.28 | 2 | 1 | 0 | 12.6 |
| 2 | 4x10<sup>6</sup> | 0.27 | unenriched | 66 | 10 | 1,478 | 1.19 | 0 | 0 | 0 | 0.0 |
| 2 | 4x10<sup>7</sup> | 2.02 | enriched | 78 | 40 | 11,005 | 7.7 | 13 | 6 | 6 | 98.6 |
| 2 | 4x10<sup>7</sup> | 2.02 | unenriched | 84 | 40 | 10,338 | 10.71 | 2 | 0 | 0 | 0.0 |
| 3 | 0 | 0.07 | enriched | 78 | 10 | 94 | 0 | 0 | 0 | 0 | 0.0 |
| 3 | 0 | 0.07 | unenriched | 66 | 10 | 349 | 0.02 | 0 | 0 | 0 | 0.0 |
| 3 | 4x10<sup>6</sup> | 0.69 | enriched | 102 | 10 | 1,442 | 2.01 | 2 | 1 | 3 | 17.2 |
| 3 | 4x10<sup>6</sup> | 0.69 | unenriched | 102 | 10 | 4,763 | 5.18 | 1 | 0 | 0 | 0.0 |
| 3 | 4x10<sup>7</sup> | 3.70 | enriched | 108 | 40 | 20,639 | 14.91 | 40 | 17 | 13 | 95.3 |
| 3 | 4x10<sup>7</sup> | 3.70 | unenriched | 108 | 40 | 18,869 | 24.45 | 0 | 0 | 1 | 13.9 |

We sought to determine how specific to *Enterobacterales* bacteria our tRNA guides would be. Guide sequences were aligned to all reference genomes of the top 391 most observed GTDB species clusters in human gut samples according to recent metagenomic sequencing studies (**Supp.** Fig 2) (see **Methods**) (Almeida et al. 2020). Overall, we found our tRNA guides to be highly specific to *Enterobacterales* – 33/36 guide sites were conserved in one or zero species, and the remaining 3 guide sites were conserved in a maximum of 16 species (range 5-16, **Supp.** Fig 2).

To enrich for AMR genes, we designed a guide pair to target the *intI1* integrase gene, a part of the class 1 integron that mobilises resistance genes and is commonly colocalised on plasmids with additional resistance determinants (see **Methods**) (White et al. 2001). We also designed four guide pairs to target highly conserved regions of the *bla*_IMP_, *bla*_OXA_ and *bla*_CTX-M_ extended-spectrum beta-lactamase (ESBL)/carbapenemase genes (see **Methods**). We prioritised targeting the alleles *bla*_IMP-4_, *bla*_OXA-48_, *bla*_CTX-M-14_ and *bla*_CTX-M-15_ as these are among the most commonly observed across Australian hospitals (Aslan and Paterson 2024). However, we expected these guides to also enrich for other allelic variants. To determine this, we calculated exact matches between our guides and each allele present in the CARD database (see **Methods**) (Alcock et al. 2020). Guide conservation varied according to allele variation in the AMR gene. For the less diverse genes *bla*_CTX-M_ and *bla*_IMP_, the guide sequences were conserved in in 77.4% (185/232) and 40.2% (33/82) of alleles respectively (**Supp Fig 3 & 4**). The more diverse *bla*_OXA_ had exact matches to the guide sequences in just 3.9% (36/912) alleles but was highly conserved in the *bla*_OXA-48_-like group of alleles known to confer carbapenem resistance (**Supp Fig 5**) (Boyd et al. 2022).

**Figure 3.**
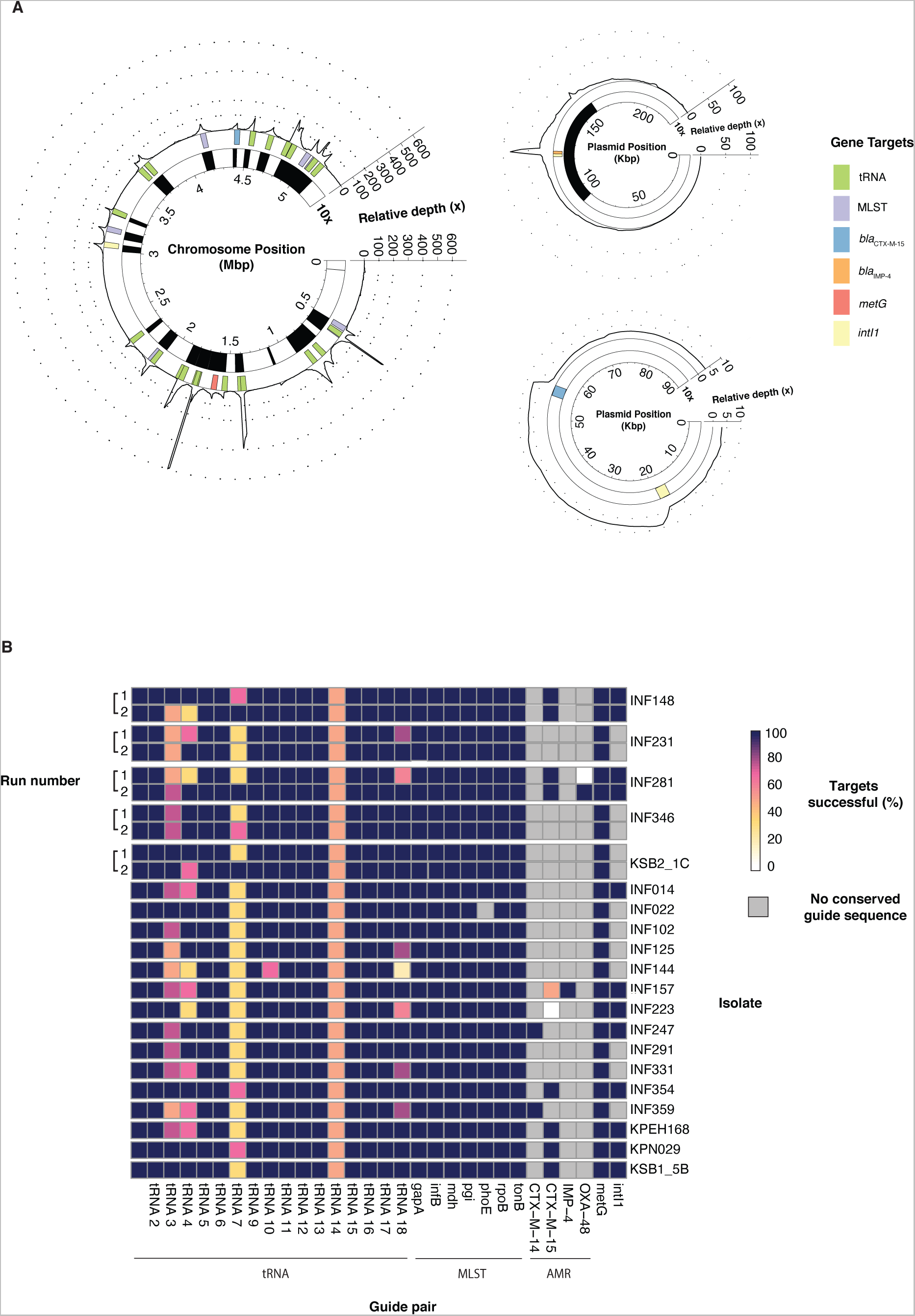
Guide pair performance in CRISPR-Cas9 enriched libraries of KpSC isolates. **A)** Sequencing depth of all target contigs in example *K. pneumoniae* isolate INF157 following CRISPR-Cas9 enrichment. Gene target locations are shown as coloured rectangles across the genome. Depth is shown relative to median depth of offtarget alignments. The inner bar denoted as ‘10x’ shows regions where relative depth is greater than 10 (shown in black). Genbank accessions are of the target contigs are CP024528.1, CP024529.1 and CP024531.1. **B)** Summary of guide performance across all 20 KpSC isolates. A successful target is defined as when the number of on-target reads is equal to or greater than 10x median depth of off-target reads. Run 1 refers to the initial sequencing with all 20 isolates, while run 2 refers to a repeat validation run on five randomly selected isolates.

**Figure 4.**
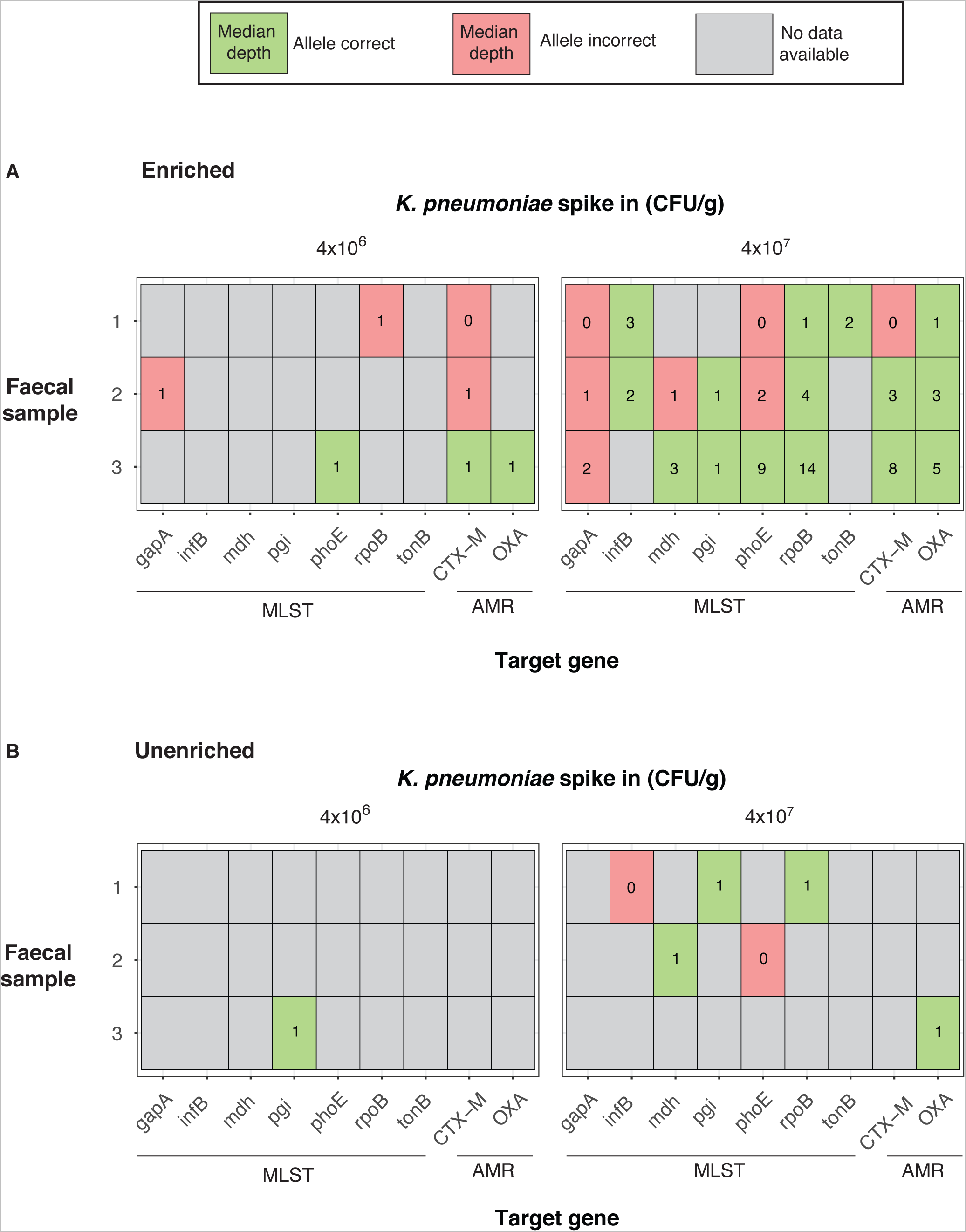
Allele call accuracy of target AMR and MLST genes following enriched and unenriched sequencing of human faecal samples spiked with *K. pneumoniae* strain INF298 (Genbank: GCA_904864465.1) at 4x10^6^ - 4x10^7^ CFU/g. Median depth was calculated from the total number of reads aligning to the target gene. **A)** Allele call accuracy following CRISPR-Cas9 enriched sequencing. **B)** Allele accuracy following unenriched sequencing.

**Figure 5.**
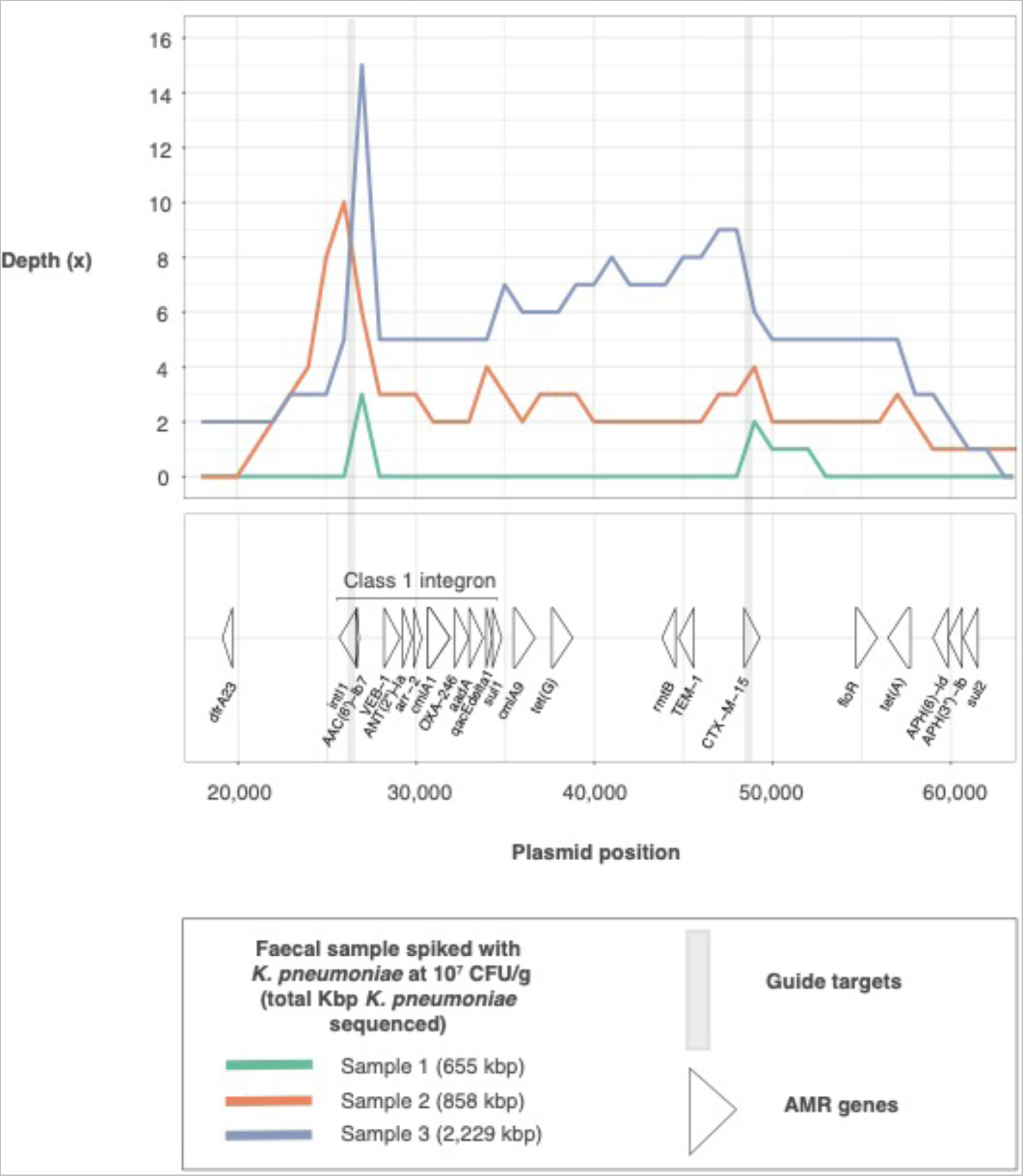
Depth sequencing of *K. pneumoniae* strain INF298 plasmid (Genbank: CP110595.1) following sequencing of human three human faecal samples spiked with INF298 at 4x10^7^ CFU/g. The bottom panel is labelled with the regions of all AMR genes on the plasmid, with the commonly observed class 1 integron labelled.

Guide conservation rate increased when focusing on clinically important mobile carbapenemase/ESBL alleles (**Supp. Table 3**, see **Methods**). We found that guides targeted 84.6% (22/26), 50% (5/10) and 63.6% (7/11) of mobile carbapenemase/ESBL alleles in *bla*_CTX-M_*, bla*_IMP_, and *bla*_OXA_ respectively. Alleles not targeted by guides were typically rarer - after adjusting for how frequently these alleles are observed in publicly available genomes, our guides were generally expected to target at least 90% of publicly available genomes possessing mobile carbapenemase/ESBL alleles (**Supp. Table 3)**. These findings suggest that although our guides were not consistently conserved in every allele of every beta-lactamase family, they likely have a high rate of enrichment in mobile carbapenemase/ESBL alleles frequently observed in clinical settings.

Finally, we designed guides to target all seven *K. pneumoniae* MLST genes, as well as the *metG* gene proximal to the K locus in *K. pneumoniae*, to enable finer-scale typing. We aligned our MLST guides to 11,446 *K. pneumoniae* publicly available genomes collected from 99 different countries over the last 100 years (see **Methods**) (Lam et al. 2021). Each guide pair matched to >99.6% of total genomes, with 89.9% (98/109) of commonly observed STs containing a guide-matching sequence in all strains (**Supp Figure 6**). Our final guide pool consisted of 31 pairs of guides (62 total) (**Supp. Table 1**).

**Figure 6.**
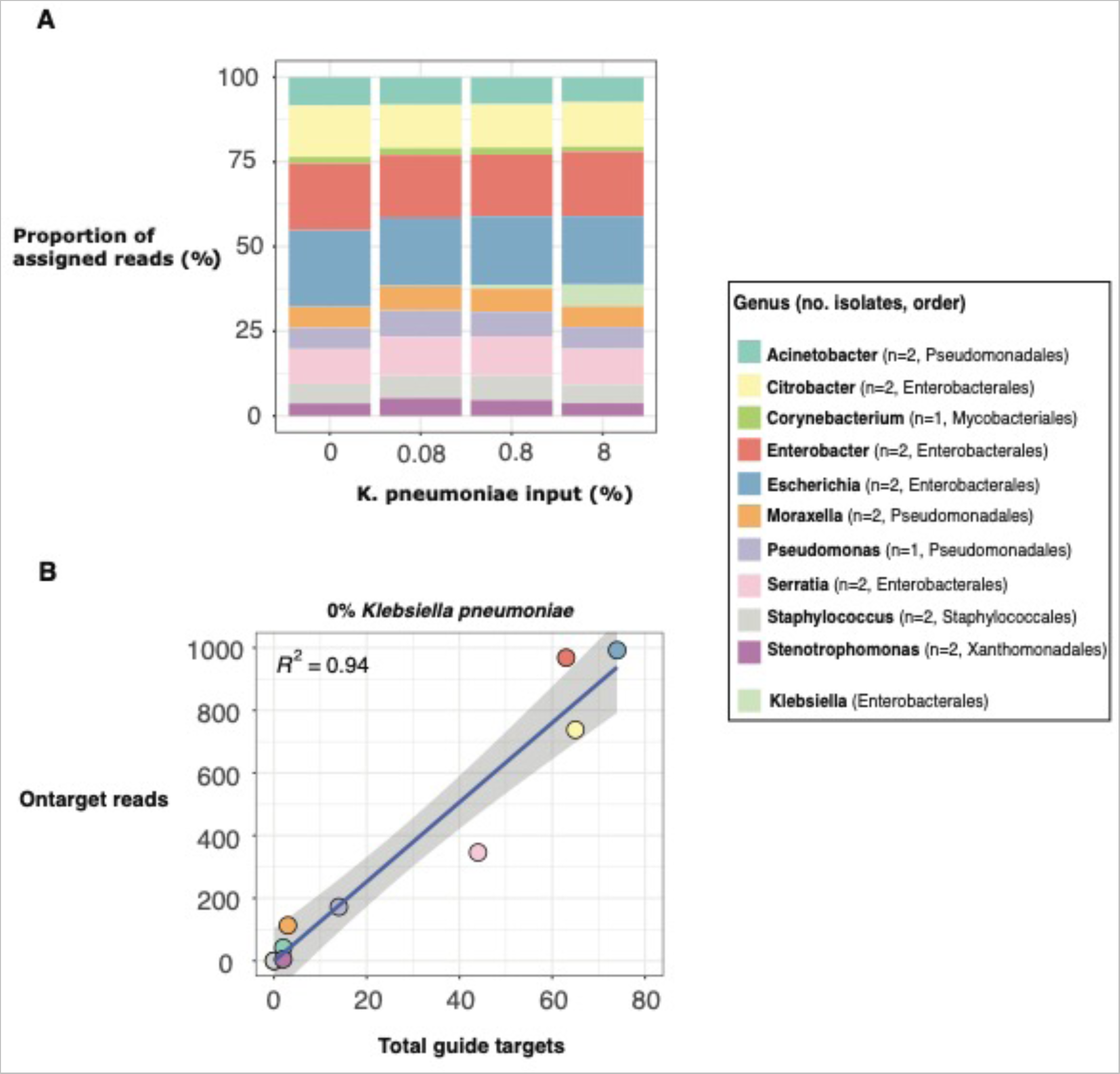
Relationship between guide conservation and enrichment performance following CRISPR-Cas9 enriched sequencing of a mock microbial community with varying amount of *K. pneumoniae* strain INF298 (Genbank: GCA_904864465.1) DNA. A) Taxonomic distribution of sequencing output based on an alignment-based approach (see **Methods**). **B)** Linear regression analysis between the total guide targets and total on-target reads for isolates of a given genus following sequencing of the mock community with no *K. pneumoniae* included.

### CRISPR-Cas9 guides consistently enriched for target sequences in *Enterobacterales* **isolates**

To assess guide performance we extracted DNA from 20 KpSC isolates, each with a publicly available completed genome and unique MLST and AMR gene profiles (**Supp. Table 4**). Nine isolates possessed an ESBL gene, and two possessed a carbapenemase gene. We performed CRISPR-Cas9 enrichment using our pool of 62 guides followed by multiplexed ONT sequencing (see **Methods**). Reads were defined as on-target if they started within 50 bp of a target site, and a guide was considered successful if the number of on-target reads was 10 times more than the background read depth estimated from off-target reads (see **Methods**).

On-target reads constituted a median of 86.7% (IQR=84.1%, 89.2%) of aligned reads across all isolates, and a median of 90.2% (IQR=86.2%, 93.3%) of conserved guide targets per isolate were successful. Successful enrichment sites were characterised by large spikes in depth surrounding target genes, with minimal sequencing in untargeted regions (**Figure 3A, Supp.** Figure 7). A median of 30.0% (IQR=24.1%, 33.5%) of each genome was recovered at above 10x depth relative to background depth, highlighting that targeting tRNA genes and using long reads can capture almost a third of the whole genome (**Supp. Tables 5-6**). All guide pairs with expected matches generated at least one successful enrichment target site in all but two isolates (*bla*_OXA-48_ in INF281 and *bla*_CTX-M-15_ in INF223, **Figure 3B**). We replicated this experiment on five randomly selected isolates, and found similar enrichment performance, this time with a successful enrichment site for *bla*_OXA-48_ in INF281 (**Run 2, Figure 3B**).

To assess the capacity of CRISPR-enriched ONT data to determine ST and AMR allelic variants in these isolates, we used on-target reads to correct random draft alleles of our target MLST and AMR genes (see **Methods**). We found that 10 reads were sufficient to produce a correct allele call in 81.8% (112/137) of cases (**Supp.** Figure 8). Accuracy continued to rise as depth increased, with 93.5% (87/93) of alleles called correctly at 50x and 97.6% (40/41) correct at 100x depth. Overall, 80% (8/10) of isolates with at least 30 on-target reads at each locus had a correct allele call in all seven MLST loci. Allele-level calls of AMR genes was comparably lower using this method, in part due to longer allele-encoding regions overall and a tendency for the majority of reads to travel from the cleavage site in a direction suboptimal to generating maximal coverage of the gene (**Supp.** Figure 9). Despite this, 7/9 isolates generated correct AMR alleles at nearly all depths. We were unable to reliably differentiate chromosomal and plasmid *bla*_CTX-M-15_ sequences following enrichment, as the surrounding ∼6 kbp of chromosomal *bla*_CTX-M-15_ genes in tested isolates showed high similarity to known plasmid sequences (see **Methods**).

To determine whether our guide pair targeting the *intI1* integron integrase gene could identify co-localised AMR genes, we used an assembly-based approach (see **Methods**). We were able to detect over half of the AMR genes present on 70% (7/10) of *intI1*-carrying plasmids by targeting either *intI1* or *intI1* plus one of our other AMR gene guides, highlighting the value of combining resistance-associated targets with long reads (**Supp. Table 7, Supp Figure 10**). Following a similar principle, we found that assembling on-target reads from the *metG* guide pair (the highly conserved gene upstream of the polysaccharide capsule-encoding K locus (see **Methods**) (Wyres et al. 2016)) allowed us to generate a correct K locus call in 60% (12/20) of *K. pneumoniae* isolates (**Supp. Table 8, Supp.** Figure 11). The relatively low success of K locus calling was due to the distance between the K locus and *metG* (∼57 kbp), which was the closest conserved sequence available to target.

Despite the success of the paired guide pool (n=62), it was not clear whether guide pairing was necessary for successful enrichment. To test this, we performed two additional enrichments using a single member of each pair rather than both (n=31). Enrichment performance was lower across all metrics, with a median of 84.4% (IQR=80.9%, 89.6%) of aligned reads coming from on-target regions and just 76.2% (IQR=74.1%, 79.0%) of targets successful per isolate. These findings were validated with a smaller repeat experiment on five of the original 20 isolates, with individual guide performance also varying in the same isolate across repeat runs (**Supp.** Figure 12**, Supp. Table 9**). For example, tRNA guides 3.1 and 2.2 were unsuccessful in all five isolates in the first run but successful in most isolates in the repeat run. Meanwhile, 4.2 and 7.2 were successful in the first run but were mostly unsuccessful in the repeat run. Overall, we found paired guides significantly outperformed unpaired guides by total targets successful (X^2^(1, N = 3482) = 65.9, *p* = 6.88 x 10^-16^) and number of libraries where every conserved guide had a successful enrichment site (X^2^(1, N = 79) = 51.1, *p* = 5.29 x 10^-12^) (**Supp. Table 10**).

By designing guides to target tRNA genes that are highly conserved in *Enterobacterales,* we aimed to enrich for several common pathogens in addition to *K. pneumoniae.* To validate this, we tested tRNA guides in a selection of *Enterobacterales* isolates not in the *Klebsiella pneumoniae* species complex. Enrichment results were similarly successful to those observed in *K. pneumoniae* isolates, with 68.4%-94.7% (26-36/38) of tRNA guides having a conserved sequence in each isolate, and each guide generating at least one successful enrichment site in isolates with a conserved sequence (**Supp. Tables 11-12**).

### CRISPR-Cas9 enrichment improved characterisation of *K. pneumoniae* from human faecal samples compared to unenriched metagenomic sequencing

We validated our CRISPR-Cas9 enrichment guides and methodology in complex patient samples, as this is their intended use-case. We spiked *K. pneumoniae* strain INF298 cells into three human faecal samples at two abundances (4x10^6^ and 4x10^7^ CFU/g, see **Methods**). We performed qPCR on all samples to determine baseline *K. pneumoniae* DNA abundance and confirm spike-ins were successful (see **Methods)**. Unspiked samples were estimated to contain 0.01-0.1% abundance of native *K. pneumoniae,* while spiked samples were estimated at 0.3-3.7% abundance (**Supp. Table 13**).

Both enriched and unenriched libraries led to detection of *K. pneumoniae* in all aliquots, typically with comparable amounts of *K. pneumoniae-*classified bases sequenced and coverage of the spiked strain genome (**Table 1**). Enriched libraries generated more *K. pneumoniae* reads aligning to MLST genes than unenriched libraries in every case, with several unenriched libraries not containing any MLST reads at all. After polishing draft sequences of MLST genes with aligned reads, enriched libraries of 4x10^7^ CFU/g spiked aliquots generated correct allele calls in 47.6% (10/21) of loci compared to 14.3% (3/21) in unenriched libraries. 4x10^6^ CFU/g appeared to be beneath the limit of consistent MLST characterisation, with correct calls in just 4.8% (1/21) loci across enriched and unenriched libraries (**Figure 4**). CRISPR-Cas9 enrichment was even more effective at picking up target AMR genes, with 55 reads aligning to AMR genes across enriched libraries compared to one read across all unenriched libraries (**Table 1**). This led to seven correct AMR allele calls in enriched libraries vs one correct AMR call in unenriched libraries (**Figure 4**). For 10/12 spiked aliquots, enrichment led to detection of target AMR genes within the first 10 hours of sequencing, with two aliquots (samples 2 and 3, 10^7^ CFU/g) obtaining detection of target AMR genes within the first hour (**Supp. Table 14, Supp.** Figure 13). Cas9 cleavage and sequencing usually led to reads in both directions from the cleavage site (**Supp.** Figure 14A**, D**) but occasionally all on-target reads travelled in a single direction, leading to poor coverage and an incorrect allele call (**Supp.** Figure 14C).

Enriched runs were able to characterise the class 1 integron present on a plasmid of the spiked strain, compared with unenriched runs which generated zero *Klebsiella-*classified reads aligning to the integron, except in one aliquot (Sample 3 10^7^ CFU/g, **Table 1**). *Klebsiella-*classified reads from spiked, enriched aliquots generated 4.5-98.6% (median 15.5%) coverage of the class 1 integron, with large spikes in depth at *intI1* (**Table 1**, **Figure 5**). While enrichment runs generated two and three reads aligning to the *metG* target gene in samples 2 and 3 spiked with 10^7^ CFU/g respectively, no *Klebsiella-*classified reads aligned to the INF298 K locus in any enriched aliquots. This is likely due to the *metG* gene being 57,219 bp away from the start of the K locus in this strain, requiring extremely long reads for effective characterisation. Unenriched aliquots generated no *Klebsiella-*classified reads aligning to the K locus except for the 4x10^7^ CFU/g spike ins, which generated one aligned read each. These reads provided limited information about the locus, with coverages of 2.4%, 7.8% and 21.3% for samples 1-3 respectively.

### Enrichment performance is highly correlated with guide conservation

To better understand how enrichment of *K. pneumoniae* was affected by guide conservation in non-target species present in a complex sample, we generated four replicate mock microbial community mixtures consisting of DNA from 11 different bacterial species at equimolar amounts, with *K. pneumoniae* strain INF298 DNA spiked in at 0%, 0.08%, 0.8%, and 8% abundance (see **Methods**). Isolates in the mock community showed varying amounts of guide conservation, ranging from 0 – 46 conserved sequences (**Supp. Table 15**). Following CRISPR-Cas9 enrichment and sequencing, we classified reads as originating from a given species using an alignment-based approach (see **Methods**). We then performed similar characterisations of the INF298 strain as in the faecal samples, using reads assigned to *K. pneumoniae*. We found trends to be very similar, with limited enrichment of *K. pneumoniae*-assigned reads (normalised to input abundance), but with high output of data from target genes (**Supp. Table 16, Supp.** Figure 15). While MLST, AMR and K locus calls were inconsistent at the 0.08-0.8% input abundance, as expected nearly all alleles were correct following enrichment and sequencing of the mixture spiked with 8% *K. pneumoniae* (Supp. Table 16).

As the mock microbial mixture was comprised of completed genomes, we could determine the relationship between guide conservation and performance (see **Methods**). We found the number of conserved guide targets to be highly predictive (R^2^ 0.90-0.94) of the amount of on-target sequence data assigned to each genus (**Figure 6**, **Supp.** Figure 16**, Supp. Tables 15, 17**).

Similarly, our faecal sequence data included on-target reads generated from taxa other than the spiked *K. pneumoniae* strain. As expected, the majority of tRNA reads were classified as originating from Proteobacteria genera such as *Klebsiella* and *Escherichia* (**Supp.** Figure 17). However, 27 reads were classified to the phylum of Bacteroidota, all of which aligned to tRNA-Gly. Guide pair three (targeting tRNA-Gly) was predicted to be conserved in many Bacteroidota species according to our initial conservation analyses, clarifying that these non-Proteobacteria tRNA sequences were in fact a by-product of our guide design rather than off-target cleavage and enrichment (**Supp.** Figure 2**, Supp.** Figure 17).

## Discussion

By using CRISPR-Cas9 enrichment to target highly conserved tRNA genes, we selectively enriched and sequenced large sections of *K. pneumoniae* and other *Enterobacterales* pathogen genomes, using DNA extracted from pure isolates, artificial isolate mixtures and human faecal samples (**Figures 2-3**, **Table 1**, **Figure 6**). We generated allele-level MLST and AMR calls by targeting highly conserved regions of these loci (**Figures 4-5**). We found CRISPR-Cas9 enrichment can outperform traditional metagenomics by identifying low abundance MLST and AMR genes from human faecal samples, with enriched libraries often picking up genes that went completely undetected in unenriched libraries (**Table 1**). Finally, we demonstrated improved enrichment consistency when using a paired guide approach, wherein multiple guides targeted overlapping regions on opposite strands of a target gene (**Supp.** Figure 12**, Supp. Table 9**). Our study is the first bacterial application to be validated in human faecal samples, enrich for tRNA genes highly conserved across *Enterobacterales* and benchmark analysis methods for identifying allelic variants from the resulting sequence data.

Although most guides generated successful enrichment in isolates with a conserved sequence, we observed large variability in depth surrounding target regions (**Figure 3B, Supp.** Figure 7). While factors such as GC content, secondary structure formation and DNA methylation have been cited as factors that may influence Cas9 cleavage success, we find this unlikely to be applicable to our context due to varying performance on the same DNA extracts over repeated sequencing runs (**Supp. Table 9)** (Hsu et al. 2013; Wu et al. 2014; Wong et al. 2015; Doench et al. 2016; Liu et al. 2016; Jensen et al. 2017; Labun et al. 2019; Chung et al. 2020). Our results indicate that designing two guides targeting overlapping regions in the target gene can mitigate some of these differences in performance and improve enrichment consistency. However, it is unclear whether improved performance was specifically due to their overlapping target sites or simply increasing the number of guides targeting the gene. A pairing approach may be best for critically important target genes such as MLST and AMR, while one guide may be sufficient for non-essential targets. Gudie conservation was highly predictive of enrichment performance in complex samples, resulting in some enrichment of Bacteroidota sequences from a tRNA guide pair (**Supp.** Figures 2 and 17). Future studies using this guide pool to enrich for *Enterobacterales* from faecal samples may wish to exclude this guide pair to avoid unintended enrichment of non-target species – guides can be easily included and excluded as desired.

This study sought to explore the how CRISPR-enriched sequence data can be used for pathogen detection and typing. Our results suggest that presence of *K. pneumoniae* MLST loci could be a reliable indicator of the presence of this species. Furthermore, the majority of MLST allelic variant calls were correct, enabling the detection of specific lineages. Enrichment of AMR gene targets was also reliable, although a higher rate of incorrect allele calls was observed than for MLST loci (**Supp.** Figures 8-9). This is likely due to the larger size of AMR genes such as *bla*_IMP_, *bla*_OXA_ and *bla*_CTX-M_, increasing the chances of inaccuracies when generating consensus sequences. Position of the guide sequence may also play a role - while MLST genes had guides target positions outside the region that specifies an allele, the entire AMR gene is allele encoding. This leads to less consistent depth across the gene and means that no single on-target read will contain the entire gene sequence. While we were able to generate accurate MLST and AMR allele calls in many cases throughout this study using ONT’s R9.4.1 sequencing chemistry, the latest R10.4.1 ONT flowcells provide much greater read-level and consensus base-call accuracy (Sereika et al. 2022) and would thus presumably yield substantially higher accuracy even with comparable amounts of data.

Yield of target sequences could be further improved via removal of non-cleaved molecules prior to sequencing. This would allow molecules with sequencing adapters attached to move more freely into the pores of the flowcell rather than being blocked by the abundance of non-target molecules still present in the solution. Some efforts have been made to improve enrichment efficiency through endonuclease depletion of untargeted DNA fragments in eukaryotic applications (Wallace et al. 2021). However, this approach is not as useful for bacterial applications where the genetic context of a target is often unknown, as it requires enrichment sites at both ends of the target region (Wallace et al. 2021). Future studies looking to improve enrichment efficiency for the single excision approach shown here could focus on molecular methods for binding to and extracting molecules with terminal phosphate groups. Meanwhile, increasing the number of guides may help to further increase the ratio of cleaved to non-cleaved DNA fragments.

Another major benefit to our ONT CRISPR-Cas9 enrichment protocol is the reduction of computational requirements compared to deep metagenomic sequencing. While typical metagenomics can often require multiple days of sequencing to reliably detect low abundance AMR genes, our CRISPR enriched libraries detected targeted *bla*_CTX-M-15_ and *bla*_OXA-48_ genes in human faecal samples from as little as one hour of sequencing. When looking at the faecal data overall, we found CRISPR-Cas9 enrichment generated 11.3x the amount of MLST reads and 56x the amount of AMR reads while using 3.3x less storage space (16GB vs 53GB) compared to unenriched sequencing of the same samples (**Table 1**). These reductions in computational requirements could substantially improve the cost effectiveness and feasibility of sequencing directly from patient samples in clinical settings. This is particularly important in contexts where resource constraints preclude traditional metagenomic approaches and low cost ONT sequencing equipment predominates.

While this study demonstrated the improved performance of CRISPR-Cas9 enrichment over typical metagenomic approaches, there are scenarios where other enrichment methods may be more appropriate. In some contexts where there is no knowledge of the disease-causing pathogen, approaches based on background depletion such as saponin depletion, CpG methylated DNA removal and depletion by hybridisation may be more suitable. While relatively straightforward and affordable, the CRISPR-Cas9 enrichment approach also requires more planning and experimental validation than computational enrichment approaches such as ONT’s adaptive sampling. While the performance of adaptive sampling has been limited so far, it may be a more accessible method for where this development is not feasible. Meanwhile, although this study displayed highly successful enrichment of target pathogens, it was limited to testing in three human samples. Future studies may look to validate performance across a larger number of human samples and wider variety of specimen types.

Our findings indicate that CRISPR-Cas9-based enrichment shows promise for targeted long-read sequencing of bacteria from clinical samples. This approach enables rapid and culture-free surveillance screening of patient samples for problematic pathogens, including *K. pneumoniae*. The additional information provided by sequencing data could inform control strategies or identify patients colonised with high-risk strains.

## Methods

### CRISPR-Cas9 guide design

To enable detection of the widest range of MLST, beta-lactamase, *intI1* and *metG* alleles as possible, we obtained a large collection of alleles for each targeted gene to identify highly conserved sequences. For the beta-lactamase genes, this collection consisted of all alleles present in a curated version of the Comprehensive Antibiotic Resistance Database (CARD) as of May 2020 (n=912 alleles for *bla*_OXA_, n=232 for *bla*_CTX-M_, n=82 for *bla*_IMP_ (Alcock et al. 2020; Lam et al. 2021). For MLST genes, the gene sequence present in strain SGH10 was aligned using BLASTn v2.13.0 (Altschul et al. 1990) to a large collection of de-replicated publicly available KpSC genomes (n=11,446) and all full-length matches to the query were retained (Lam et al. 2021). This dataset included genomes from 99 countries collected from animal, environmental, food and human sources over the last 100 years (Lam et al. 2021). *intI1* alleles were identified by aligning the publicly available reference gene (Genbank accession CP024557.1) to the same KpSC genome collection and retaining full-length matches. *metG* alleles were identified using panaroo v1.2.2 (Tonkin-Hill et al. 2020) from a set of 328 KpSC genomes collected between 2013-2014 from the Alfred Hospital in Melbourne, Australia (Gorrie et al. 2017; Gorrie et al. 2018; Gorrie et al. 2022). The study was reviewed and approved by the Alfred Hospital Ethics Committee.

All alleles from each target gene (with the exception of *metG*, where we utilised the alignment from panaroo) were aligned using MUSCLE v3.8.31 (Edgar 2004). Highly conserved sequences were visually identified in Jalview v2.11.1 (Waterhouse et al. 2009). For MLST genes, we ensured that the conserved regions selected were outside of the MLST allele-coding region to facilitate MLST typing. All conserved regions were input into the CRISPR-Cas9 guide design tool CHOPCHOP v3 (Labun et al. 2019) using the ‘nanopore enrichment’ setting, with *Homo sapiens* hg38/GRCh38 as the background organism to minimise potential matches to human DNA. Preference was given to guides with minimal close matches to the background genome (MM1=0, MM2<3, MM3<5), %GC ranging from 40-60%, and no self-complementarity. Based on preliminary results, we hypothesised that designing two guides with conserved regions on opposite strands of target genes would be more effective than a single guide per target sequence. To test this, we designed guides with these overlapping regions for each target gene. If the ‘nanopore enrichment’ setting did not yield two guides with overlapping regions, we used the default ‘knock-out’ setting to produce a larger pool of candidate sequences. We ordered 31 guide pairs (n=62 total) guides from Integrated DNA Technologies using the Custom Alt-R® CRISPR-Cas9 guide RNA tool (**Supp. Table 1**).

### Analysis of guide conservation

To assess tRNA guide conservation within *Enterobacterales*, we first prepared a dereplicated version of *Enterobacterales* genomes in the Genome Taxonomy Database (GTDB) (release 95) (Parks et al. 2018). Each GTDB species was dereplicated with Assembly-Dereplicator v0.1.0 (Wick and Holt 2023), first using a distance threshold of 0.001, then increasing the threshold until either the number of assemblies dropped below 100 or the threshold reached 0.05 (n=11,339 total *Enterobacterales* genomes). We then aligned tRNA guides to all *Enterobacterales* genomes using bowtie2 v2.3.5.1 (Langmead and Salzberg 2012) and summarised the proportion of genomes in each genus with a perfectly conserved guide. To visualise genera in *Enterobacterales* and *Klebsiella,* a single representative genome was taken from each genera or species respectively and used as input into mashtree v1.2.0 (Lee S. Katz 2019). For conservation outside *Enterobacterales,* guides were aligned to all genomes of the top 500 most commonly observed GTDB (release 89) species in human gut samples (Almeida et al. 2020) using bowtie2 v2.3.5.1 (Langmead and Salzberg 2012). Species clusters that were unclassified (n=91), duplicated (n=5), or members of *Enterobacterales* (n=13) were excluded for a final dataset of 391 species clusters.

AMR guides were aligned to all alleles of each target gene present in the CARD database as of June 2022 using bowtie2 v2.3.5.1 (Langmead and Salzberg 2012; Alcock et al. 2020). Multiple sequence alignment and BioNJ trees of all alleles for each gene were generated in seaview (Gascuel 1997; Gouy et al. 2009). Mobile carbapenemase/ESBL alleles were defined in this study as those found in multiple *Enterobacterales* species according to CARD prevalence data. Prevalence data was then used to summarise their rate of carriage in public assemblies and the rate at which those assemblies contain conserved guide sequences. MLST guides were aligned to the previously described database of 11,446 *K. pneumoniae* genomes (Lam et al. 2021) using bowtie2 v2.3.5.1 (Langmead and Salzberg 2012). All conservation calculations were performed in R.

### Sample preparation

Bacterial isolates were grown overnight on LB agar and DNA extraction was performed using the GenFind v3 gDNA extraction kit according to standard protocol (**Supp. Table 4**) (Beckman Coulter). For the mock microbial community, we pooled equimolar amounts of genomic DNA from 18 bacterial isolates (**Supp. Table 14**). *K. pneumoniae* strain INF298 (GenBank accession GCA_904864465.1) genomic DNA was spiked into four duplicate aliquots of the community at varying relative abundance (0%, 0.08%, 0.8% and 8%).

For faecal experiments, we mixed 0.3g from each sample in 1mL of sterile 1X PBS to ensure even bacterial distribution. We then took 3 aliquots (0.1g faeces each) of each sample and spiked in 0, 4x10^5^, or 4x10^6^ CFU of *K. pneumoniae* strain INF298 cells grown overnight in Luria-Bertani broth. This resulted in 0.1g faecal aliquots with *K. pneumoniae* strain INF298 spiked in at concentrations of 0, 4x10^6^ and 4x10^7^ CFU/g; an estimated range that would typically be found in faecal samples (Huttenhower et al. 2012; Sender et al. 2016; Rothschild et al. 2018). We extracted faecal DNA using the ‘Three Peaks’ method to retain long DNA fragments for Oxford Nanopore sequencing (Quick 2019). Briefly, this involves first removing free DNA present in the sample, then enzymatic cell lysis and DNA extraction followed by bead beating and DNA extraction.

### Quantitative PCR (qPCR)

To determine *K. pneumoniae* abundance in faecal DNA extracts, qPCR was conducted using Promega’s GoTaq reagents with primers specific to *K. pneumoniae* and a standard curve of pure INF298 genomic DNA (Barbier et al. 2020). Sample, reagent, and thermocycler details can be found in **Supp. Table 13**.

### CRISPR-Cas9 enrichment and DNA sequencing

CRISPR-Cas9 enrichment was performed according to Oxford Nanopore’s Cas9 Targeted Sequencing protocol with some modifications to facilitate multiplexed libraries (**Supp. Methods 1**). Briefly, this involved preparing Cas9 ribonucleoproteins (RNPs) by combining *Streptococcus pyogenes* Cas9 Nuclease, tracrRNA and crRNAs. Genomic DNA was then dephosphorylated using calf intestinal phosphatase to prevent adapter ligation. Cas9 cleavage was induced at target sites to expose DNA terminal phosphate groups and allow for adapter ligation in these areas (**Figure 1**). Final DNA libraries were prepared using ligation kit LSK-109 and the barcoding expansion kit EXP-NBD196, sequenced on R9.4.1 MinION flowcells and basecalled using the super model of guppy v6.2.1 for isolate experiments and v7.1.4 for faecal experiments. For faecal experiments, we delayed pooling barcodes until after adapter ligation to minimise any cross-barcode leakage (**Supp. Methods 2**). Enriched and unenriched libraries of each faecal sample were run for the same duration (Ten hours for 0 CFU/g and 4x10^6^ CFU/g aliquots, 40 hours for 4x10^7^ CFU/g aliquots) using separate flowcells with comparable pore counts.

### Analysis of enrichment success

For isolate and mock community experiments, we used previously completed genomes (**Supp. Table 4, Supp. Table 15**). Guide sequences were aligned to assemblies using bowtie2 v2.5.1 with the -a parameter to identify all target regions (Langmead and Salzberg 2012). Reads were then aligned to assemblies using minimap2 v2.24 (Li 2018) with the -map-ont, -c and --secondary=no parameters. After aligning CRISPR-enriched sequence data to the completed genome of each isolate, on-target reads were defined as those with alignments starting or ending within 20bp of the conserved guide site. These alignments were most likely the result of Cas9 cleavage, adapter ligation and sequencing beginning at these sites. Successful guide sites were defined as those with ten or more on-target reads divided by the median depth of off-target (non-on-target) reads for that contig. Normalising on-target read levels to the depth of off-target reads was to account for yield differences between isolates. To visualise enrichment performance, the depth of sequencing at each position in the genome was calculated using samtools depth v1.1.7 (Danecek et al. 2021). Circular depth plots were generated using the R package circlize v0.4.10, while linear depth plots were generated in ggplot2 (Gu et al. 2014). Chi-Squared tests to compare paired and unpaired guide performance were generated in R using the chi.test function with default settings.

To assign reads to species following CRISPR-Cas9 enrichment and sequencing of the mock microbial community, we first aligned reads to each of the isolates in the mixture We then classified them as originating from a given species if the highest scoring alignments for at least 80% of positions in the read stemmed from isolates of that species. Linear regression analysis to assess the relationship between guide conservation and performance in the artificial bacterial mixture was performed using the ggpmisc package. A p value less than 0.05 was treated as statistically significant.

For faecal experiments, we performed species classification using Kraken 2 with the GTDB database release 202 as reference (Méric et al. 2019; Wood et al. 2019). To extract reads classified into taxons of interest we used read IDs from the Kraken 2’s output as input to seqtk’s subseq command (https://github.com/lh3/seqtk). To determine the number of reads aligning to target loci, we aligned reads to fasta files of the target genes using minimap2 with the -c and --secondary=no flags. To extracts reads according to sequencing duration, we sorted fastq read IDs by the start_time flag on the header line and calculated the elapsed time since the first read of the run. We then used the read IDs that were beneath a given sequencing duration as input into seqtk’s subseq command (https://github.com/lh3/seqtk).

### Characterising MLST and AMR genes using enriched sequences

We generated consensus sequences of target MLST and AMR genes by using raw reads to polish a random allele known not to match the allele present in each genome using medaka v1.5.0 (https://github.com/nanoporetech/medaka). For MLST genes, the draft sequence was a randomly chosen allele and for AMR genes we chose a random allele from the same clade as the target allele. We found that changing the draft allele within these parameters had no noticeable effect on consensus accuracy.

To determine whether we could differentiate between chromosomal and plasmid *bla*_CTX-M_ reads following CRISPR-Cas9 enrichment and sequencing of *K. pneumoniae* isolates, we classified reads using Kraken 2 (Wood et al. 2019) with a custom database of *Enterobacterales* chromosomes and plasmids (Gomi et al. 2021). The database included all fully assembled KpSC chromosomes found in NCBI and all complete *Enterobacterales* plasmids in PLSDB as of December 2023 (Galata et al. 2019).

To determine how many *intI1*-adjacent AMR genes we could identify in isolate experiments, we assembled all on-target reads from each *intI1*-containing plasmid using flye v2.9 (Kolmogorov et al. 2019) with 70,000 as the --genome-size parameter. We ran the resulting assemblies through Kleborate v2.3.2 (Lam et al. 2021) with the -r parameter and compared AMR results to those run on the plasmid from the completed assembly. For *metG* analyses, we assembled on-target *metG* reads using flye v2.9 (Kolmogorov et al. 2019) with 70,000 as the --genome-size parameter, ran Kaptive v3.0.0b (Wyres et al. 2016; Lam et al. 2022) and compared results to the completed assembly. For faecal experiments, we extracted reads that aligned to the *intI1* gene and aligned those to the June 2022 version of the CARD AMR database (Alcock et al. 2020).

## Data Access

All sequence data generated in this study have been deposited in the NCBI database under the BioProject accession PRJNA1123839.

## Acknowledgements

This research was supported by use of the Nectar Research Cloud, a collaborative Australian research platform supported by the NCRIS-funded Australian Research Data Commons (ARDC) and by award OPP1210746 to INO and KH from the Bill & Melinda Gates Foundation. INO is a Calestous Juma Science Leadership fellow supported by the Bill & Melinda Gates Foundation (INV-036234).

We also acknowledge the work of Helena Cooper, who adapted the database used for differentiating between chromosomal and plasmid sequences for our application.

## Notes

### Competing Interest Statement

The authors have declared no competing interest.

