## Supplemental Figures for "Targeted sequencing of *Enterobacterales* bacteria using CRISPR-Cas9 enrichment and Oxford Nanopore Technologies"

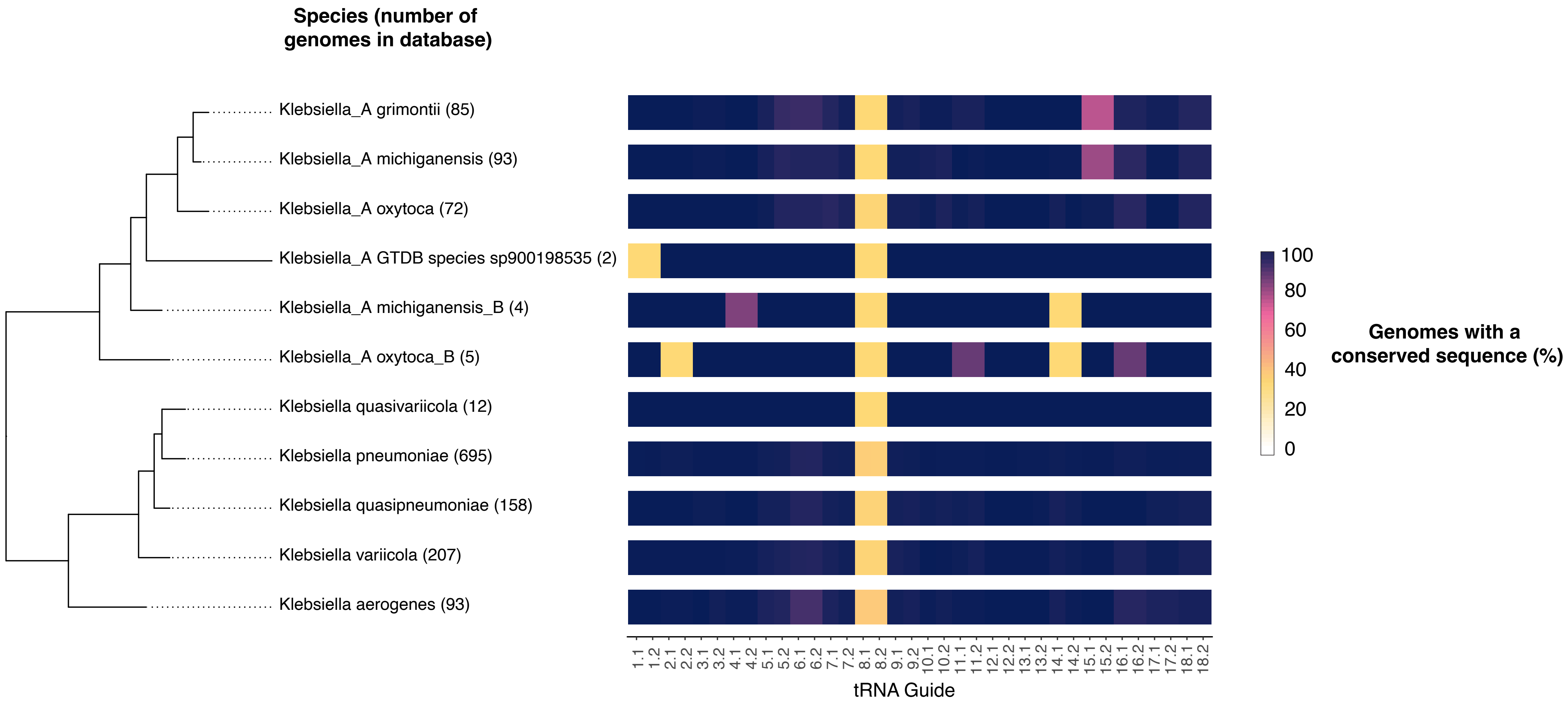


**Supplementary Figure 1 - Conservation of tRNA guides across *Klebsiella***

To the left is a neighbour-joining tree of representative genomes from all genera in GTDB r95 classified as *Klebsiella* or *Klebsiella_A* (one genome per species, n=11 species) (Parks et al. 2018; Lee S. Katz 2019). The colour spectrum of the heatmap shows the proportion of genomes matched to guide sequences in a dereplicated version of the full GTDB database for each species (n=1,426 total *Klebsiella* or *Klebsiella_A* genomes, genome count for each species shown in brackets).

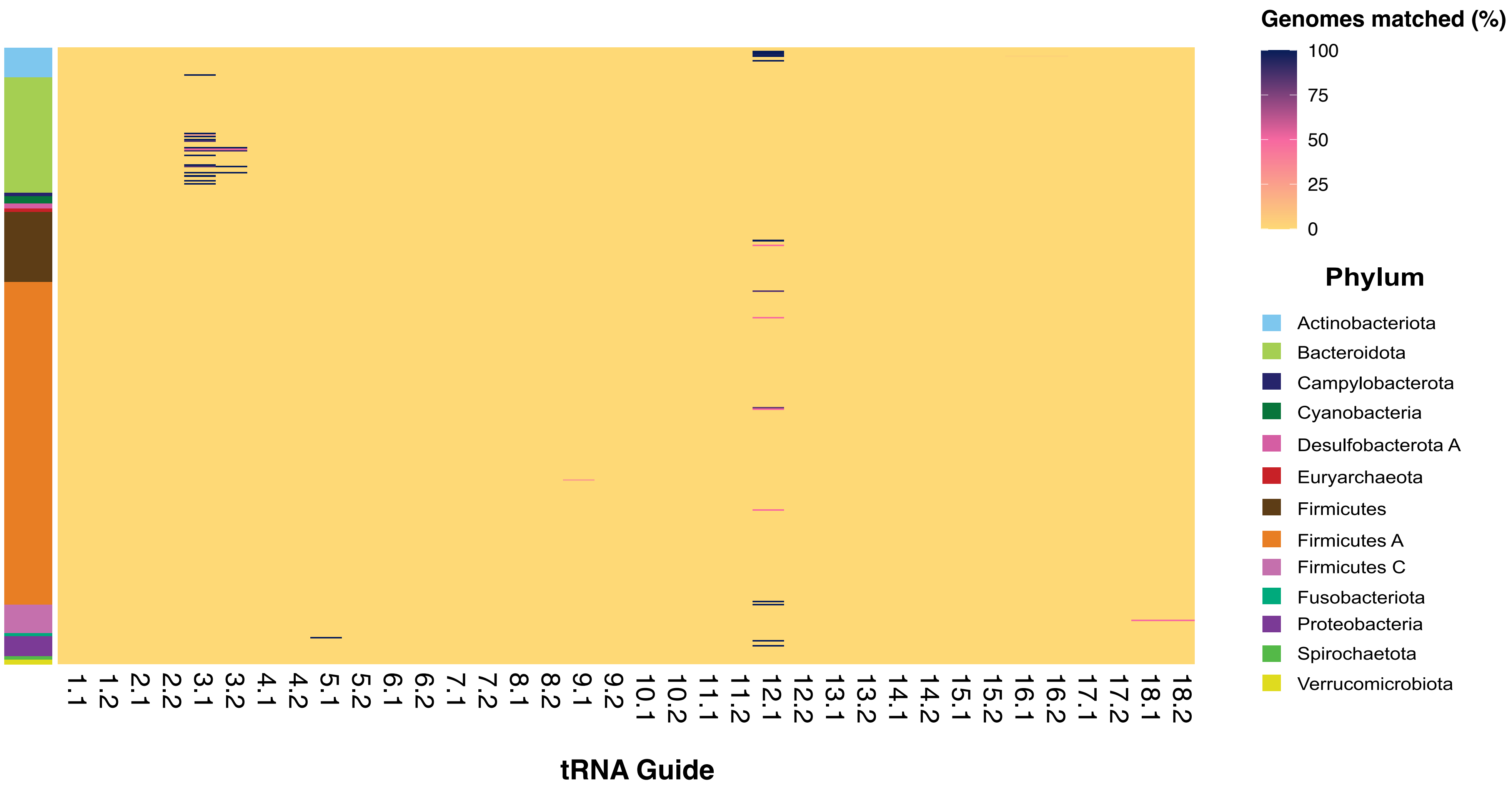


**Supplementary Figure 2 - Conservation of tRNA guides amongst commonly observed species outside of *Enterobacterales***
Rows show each of the top 391 most abundant GTDB-defined species present in the human gut (Parks et al. 2018; Almeida et al. 2020). The colour spectrum refers to the proportion of genomes assigned to a given species in the GTDB database that matched with guides. The colour bar to the left of heatmap shows GTDB-defined phylum of each species.

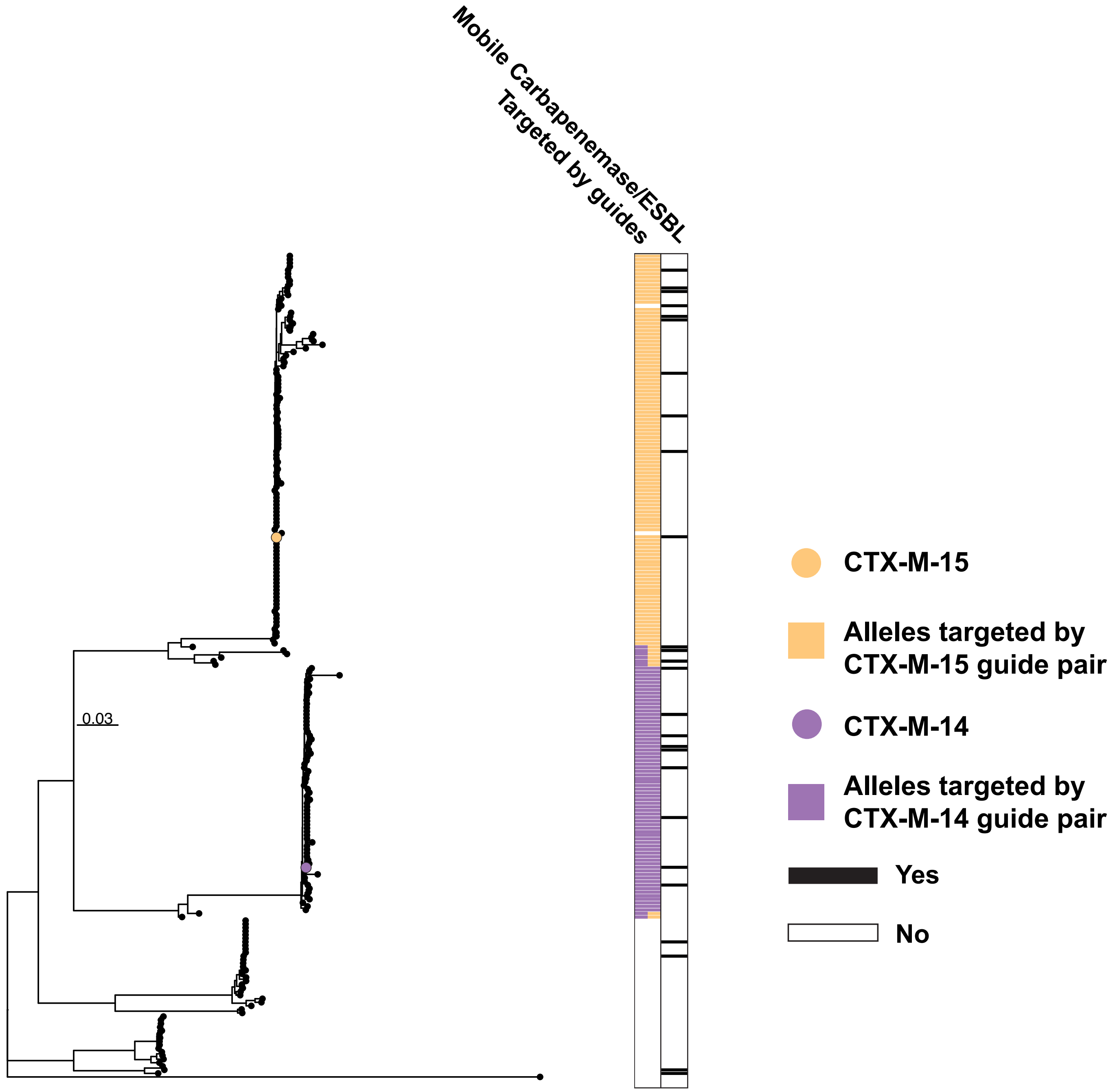


**Supplementary Figure 3 - Phylogenetic tree of all *bla*$_{\text{CTX-M}}$ alleles and those targeted by *bla*$_{\text{CTX-M}}$ guides**
Generated using BIONJ, rooted at centre (Gascuel 1997; Alcock et al. 2020; Lam et al. 2021). Each dot (node) represent a *bla*$_{\text{CTX-M}}$ allele. The nodes of primary target alleles are coloured, with each allele targeted by the two guide pairs shown as colours in the heatmap. Some alleles are targeted by both pairs- these are represented by bars with both blue and red segments. Alleles that are ESBL/carbapenemase and found in multiple *Enterobacterales* species are also denoted in column two of the heatmap. The scale bar shows the distance correlating to 2% nucleotide variation between clades.

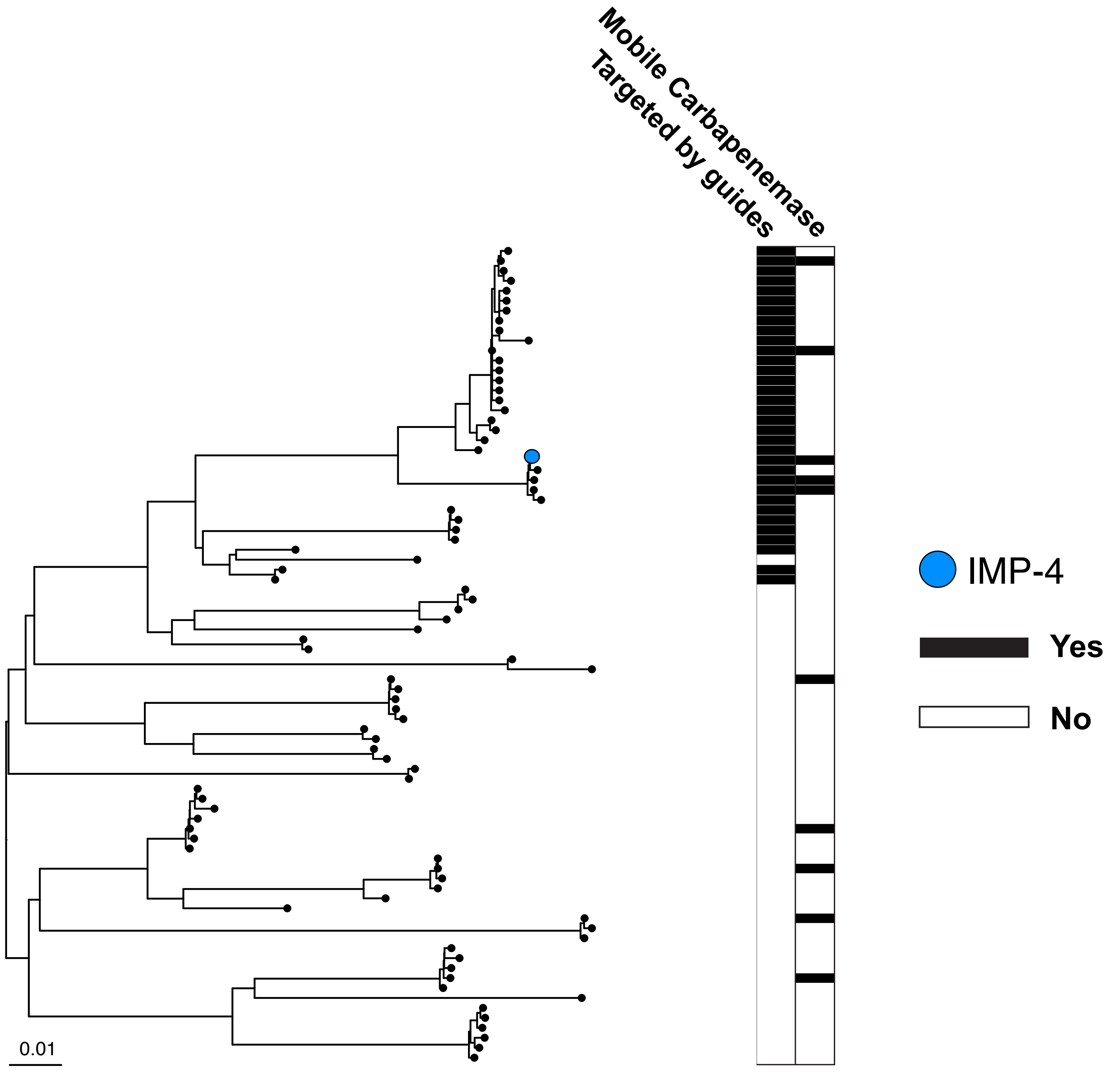


**Supplementary Figure 4 - Phylogenetic tree of all *bla*IMP alleles and those targeted by *bla*IMP guides**
Generated using BIONJ, rooted at centre (Gascuel 1997; Alcock et al. 2020; Lam et al. 2021). Each dot (node) represent a *bla*IMP allele. The node of the primary target allele is coloured in blue, with each allele targeted by the guide pair shown as black in first row of the heatmap. Alleles that are carbapenemase found in multiple *Enterobacterales* species are also denoted in the second row. The scale bar shows the distance correlating to 1% nucleotide variation between clades.

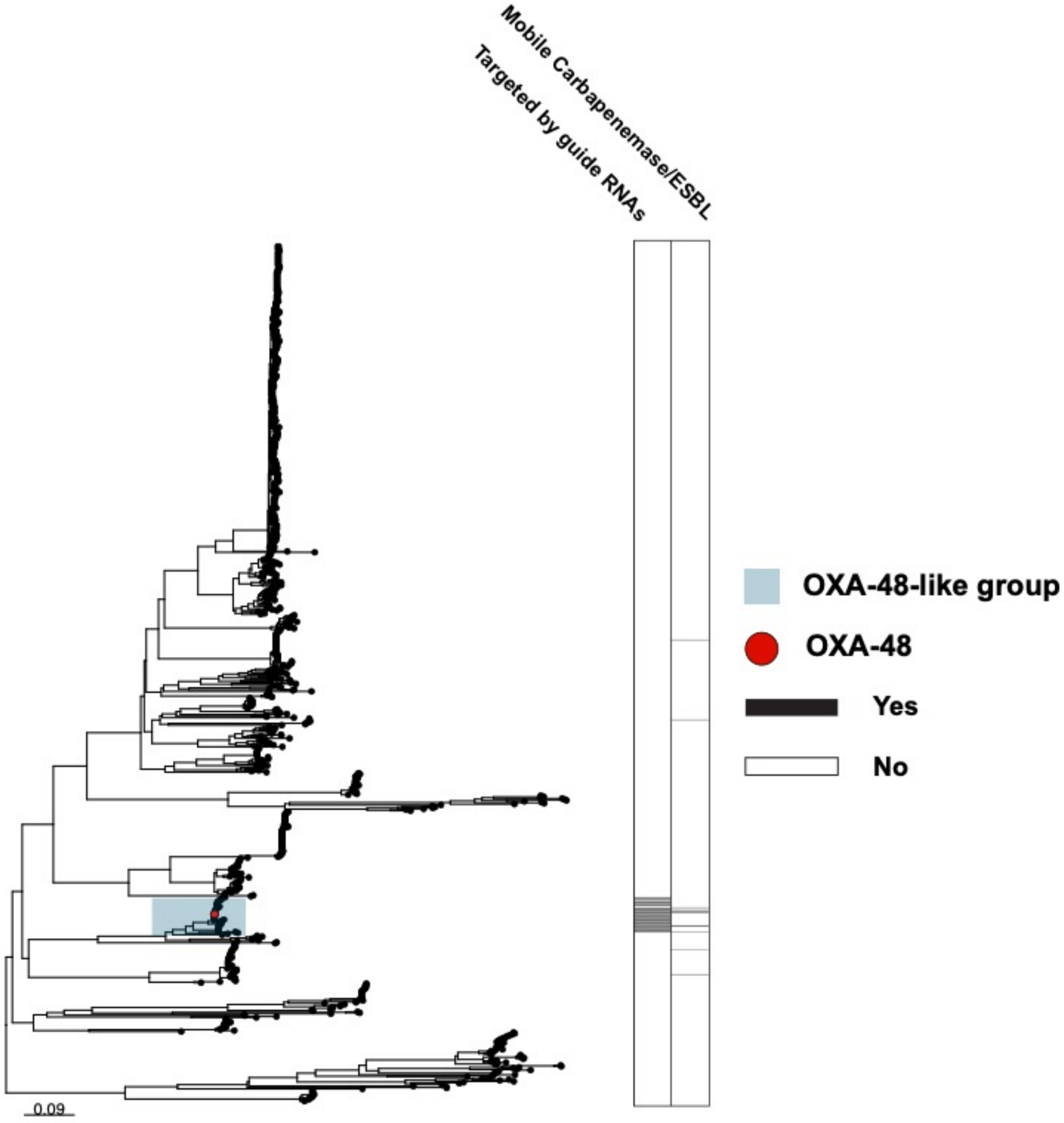


**Supplementary Figure 5 Phylogenetic tree of all *bla*$_{OXA}$ alleles and those targeted by *bla*$_{OXA}$ guides**

Generated using BIONJ, rooted at centre (Gascuel 1997; Alcock et al. 2020; Lam et al. 2021). Each dot (node) represent a *bla*$_{OXA}$ allele. The node of the primary target allele is coloured in red, with each allele targeted by the guide pair shown as black in the first row of the heatmap. The OXA-48-like group of alleles are highlighted in blue. Alleles that are carbapenemase and found in multiple *Enterobacterales* species are also denoted in the second column. The scale bar shows the distance correlating to 7% nucleotide variation between clades.

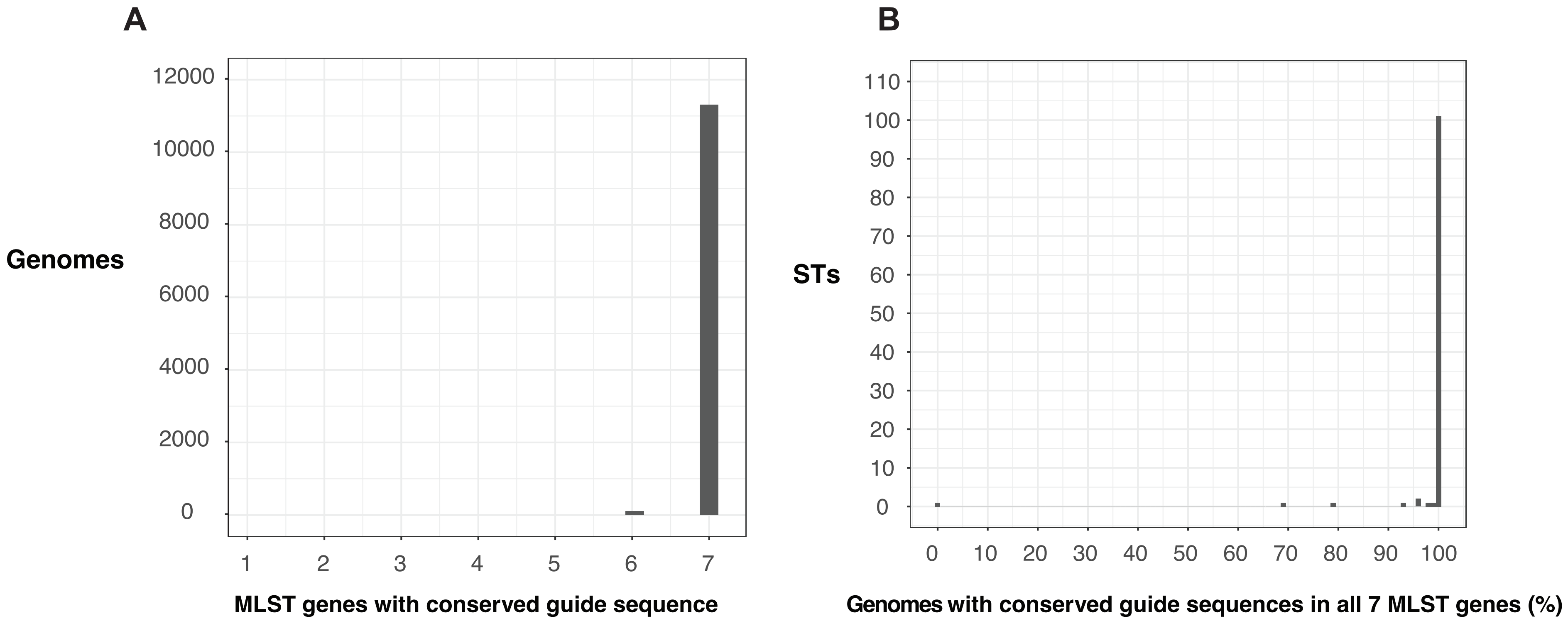


**Supplementary Figure 6 – Conservation of MLST-targeting guide pairs in 11,346 *K. pneumoniae* genomes.**

Both members of each pair must have a conserved sequence for a match to be reported. **A)** Distribution of the number of MLST genes containing conserved guide sequences for a given genome. **B)** Distribution of number of genomes, for each ST greater than 1% abundance, with a perfect match to all MLST guides.

**Supplementary Figure 7** – Sequencing depth of *Enterobacterales* isolates containing conserved guide sequences following CRISPR-Cas9 enrichment and ONT sequencing.

Contig number is shown under each isolate code.Total depth is shown as the dark blue line, with offtarget depth in light blue. Target gene regions are shown as coloured rectangles.

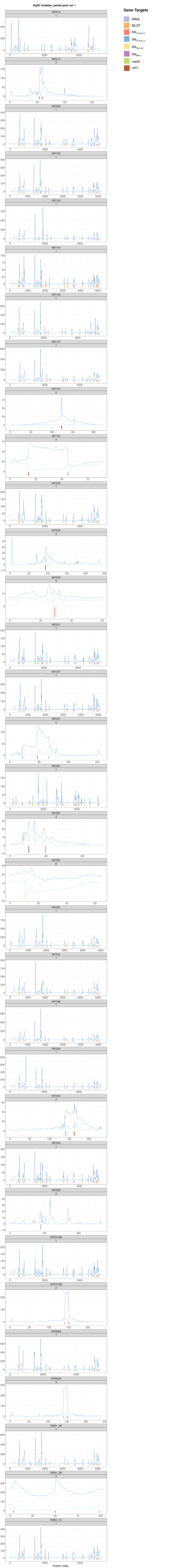

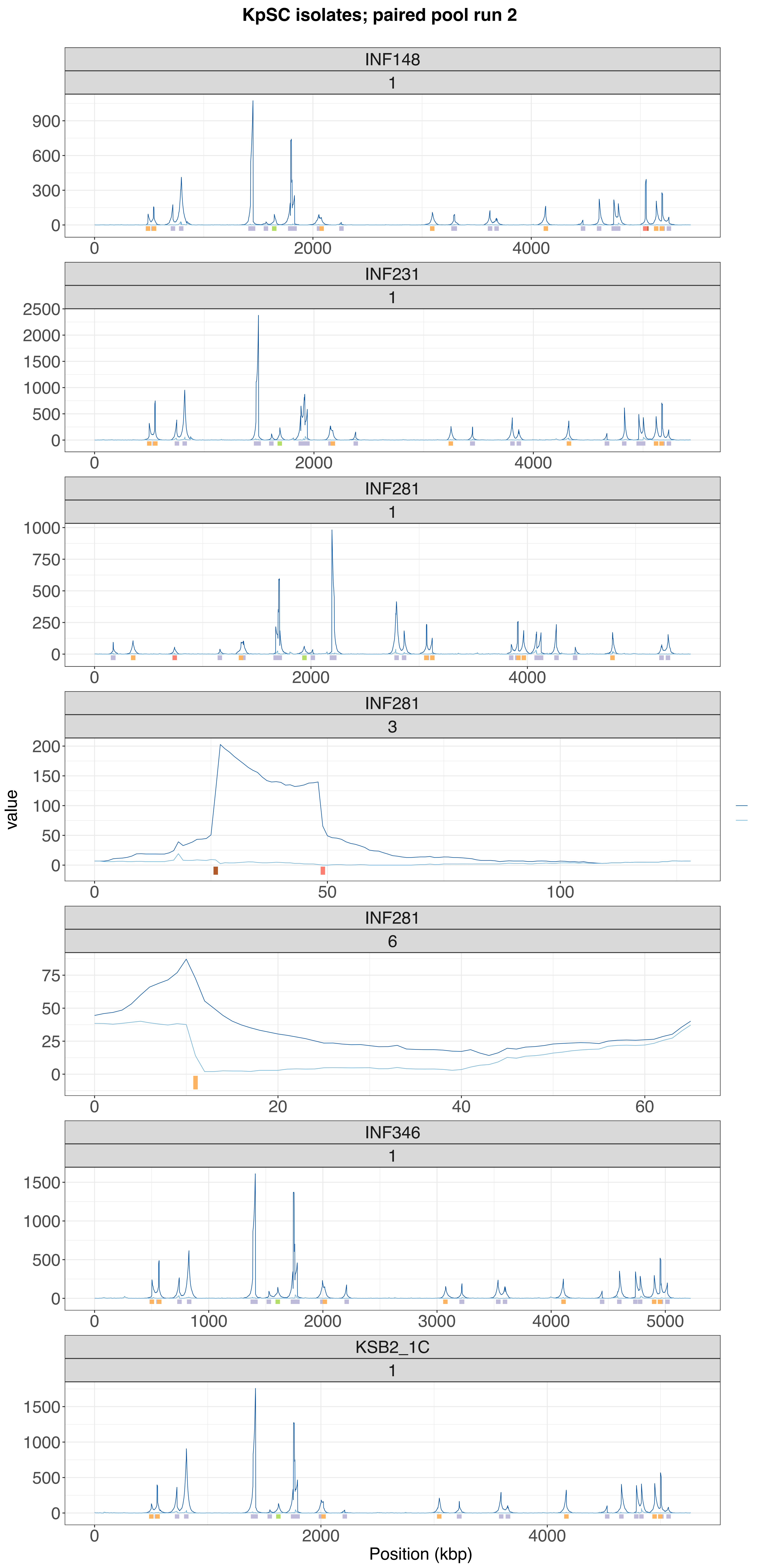
KpSC isolates; paired pool run 2
INF148
1
INF231
1
INF281
1
INF281
3
INF281
6
INF346
1
KSB2_1C
1
value
Position (kbp)

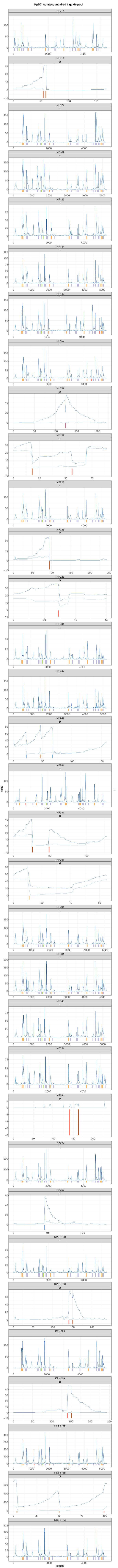
KpSC isolates; unpaired 1 guide pool
value
region

**KpSC isolates; unpaired 2 guide pool**

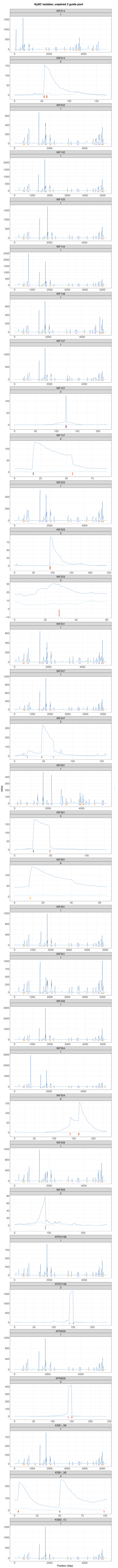

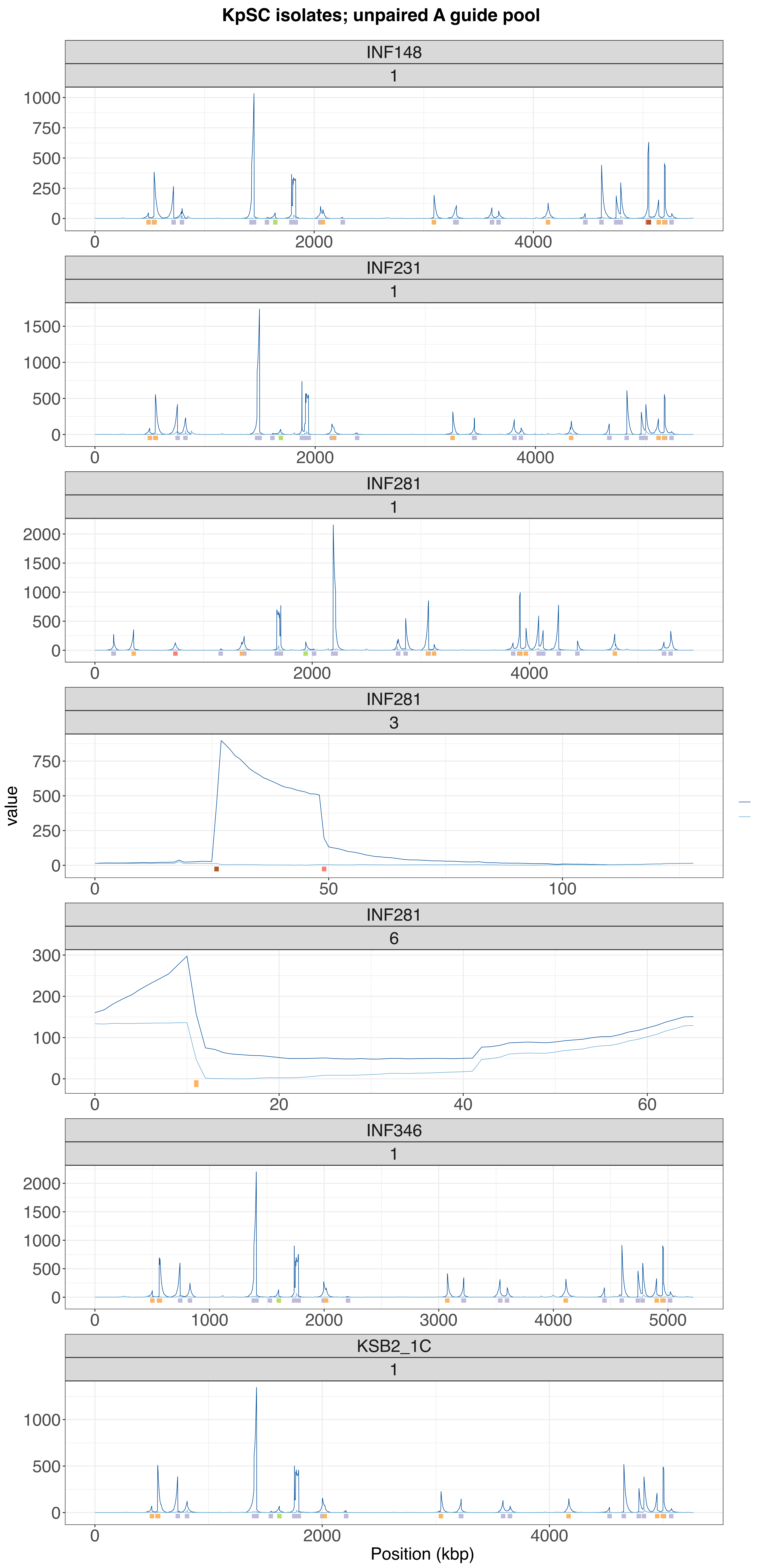

KpSC isolates; unpaired A guide pool
INF148
1
INF231
1
INF281
1
INF281
3
INF281
6
INF346
1
KSB2_1C
1
value
Position (kbp)

**KpSC isolates; unpaired B guide pool**

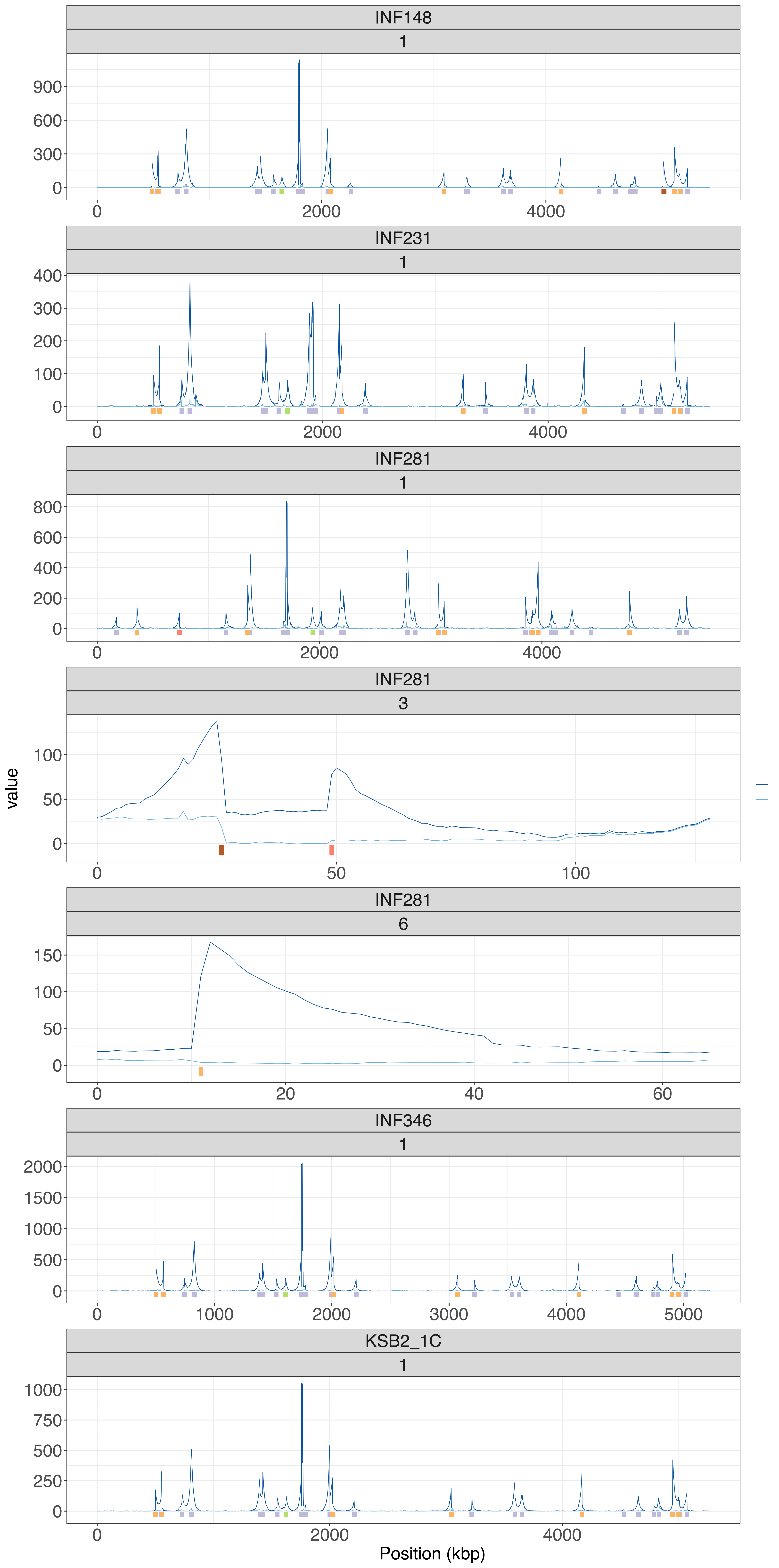

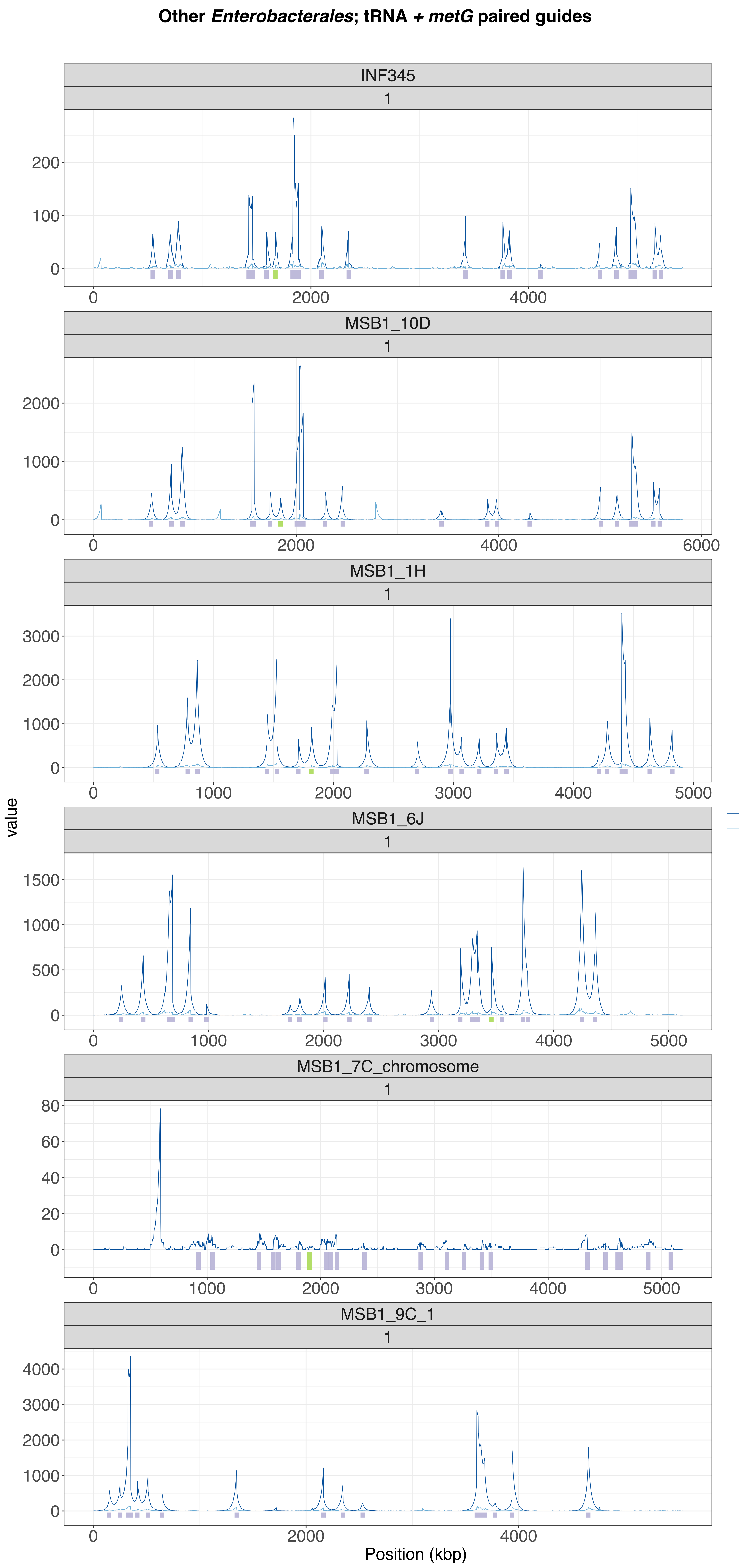
Other *Enterobacterales*; tRNA + *metG* paired guides
INF345
1
MSB1_10D
1
MSB1_1H
1
MSB1_6J
1
MSB1_7C_chromosome
1
MSB1_9C_1
1
value
Position (kbp)

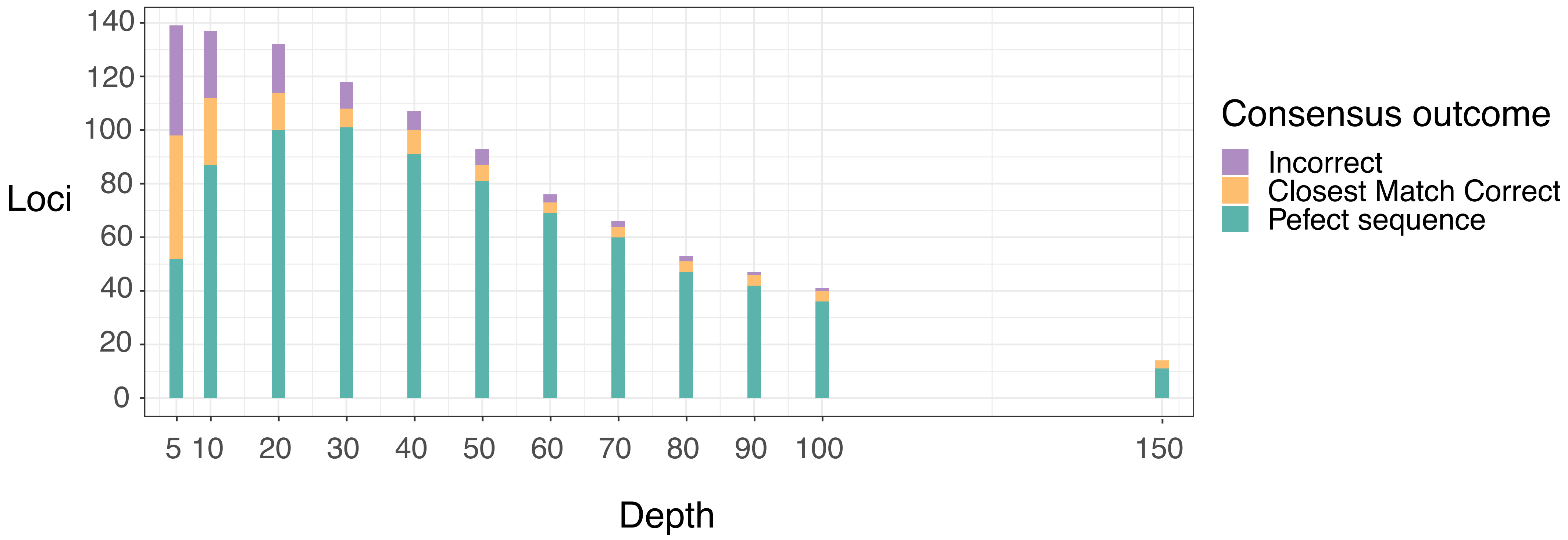


**Supplementary Figure 8 – Accuracy of MLST consensus sequences generated from CRISPR-Cas9 enriched ONT data of 20 *K. pneumoniae* isolates.**

Depth refers to the number of ontarget reads spanning the MLST-encoding region of a given MLST gene. Consensus outcomes are measured against the completed assembly for that given isolate. ‘Incorrect’ consensus sequences refer to those with errors in their sequence and with an incorrect closest allele match when aligned against the MLST database. ‘Perfect sequence’ refers to sequences where the allele is correct, and every base is correct. ‘Closest Match Correct’ refers to consensus sequences with at least one incorrect base but the top scoring allele is correct when aligned to the MLST database.

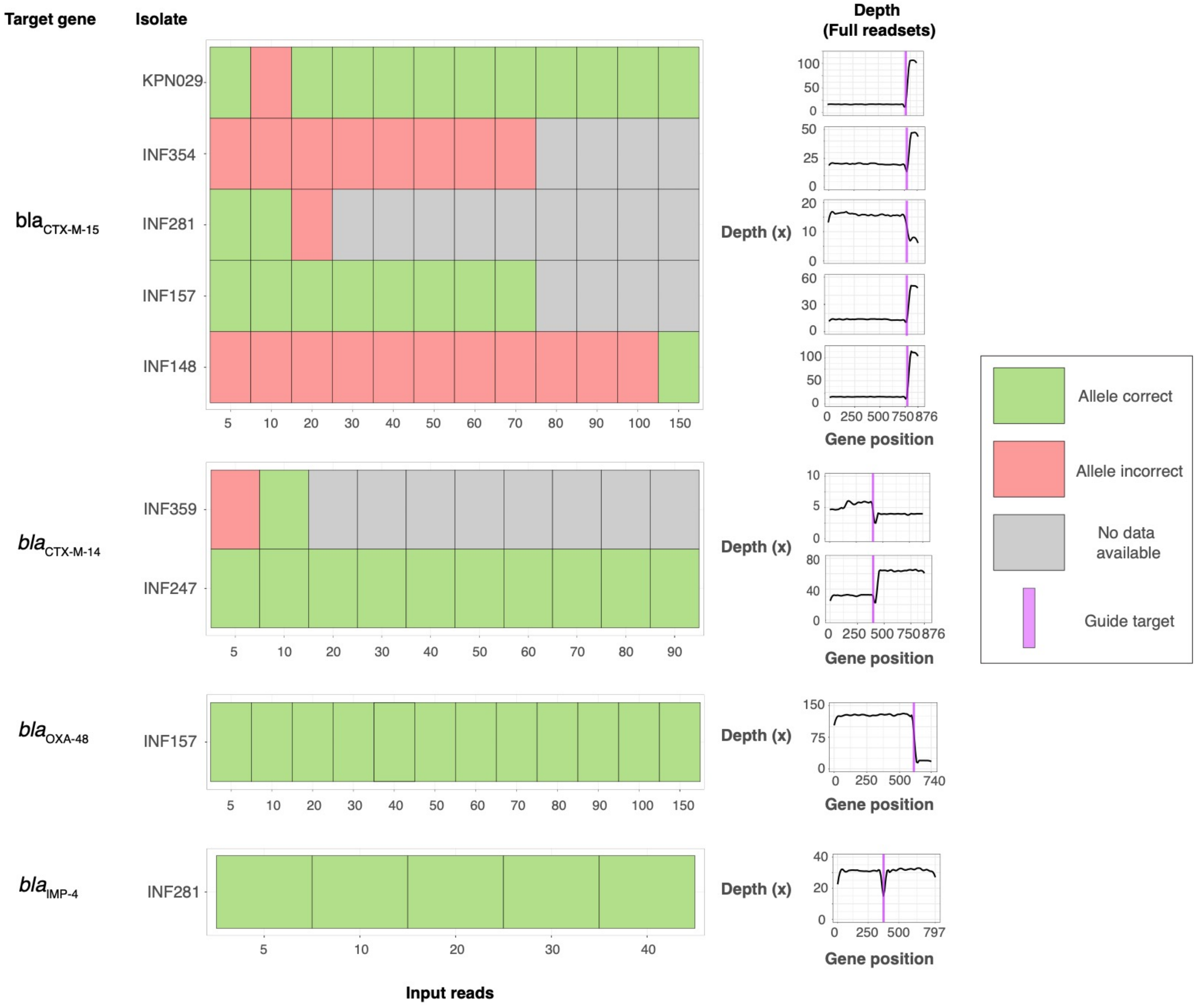


**Supplementary Figure 9 – Accuracy of AMR consensus sequences generated from CRISPR-Cas9 enriched ONT data of 20 *K. pneumoniae* isolates.**

Input reads represent random subsets of the full collection of reads aligning to the target AMR gene. Consensus outcomes are measured against the completed assembly for that given isolate. 'Incorrect' consensus sequences refer to those with errors in their sequence and with an incorrect closest allele match when aligned against the MLST database. 'Correct' refers to consensus sequences where the top scoring allele is correct when aligned to the CARD AMR database (Alcock et al. 2020).

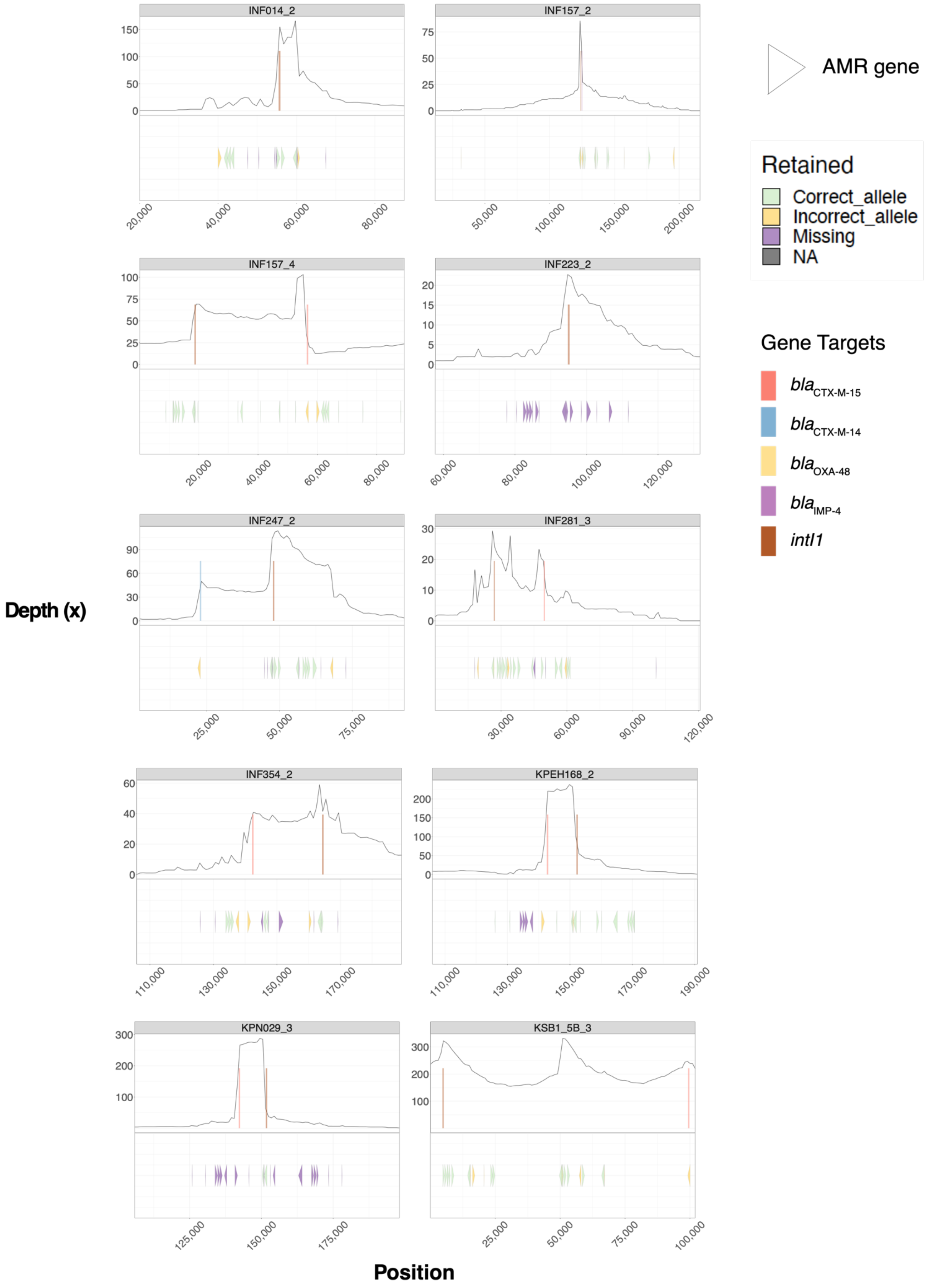


**Supplementary Figure 10 – Depth of ontarget reads aligned against *intI1*-containing plasmids across *K. pneumoniae* isolates following CRISPR-Cas9 enrichment and ONT sequencing.**
Guide target regions are shown in coloured rectangles. On the bottom panel is the regions of AMR genes on the plasmid, coloured by whether they were absent, retained with the correct allele or retained with an incorrect allele once reads were assembled.

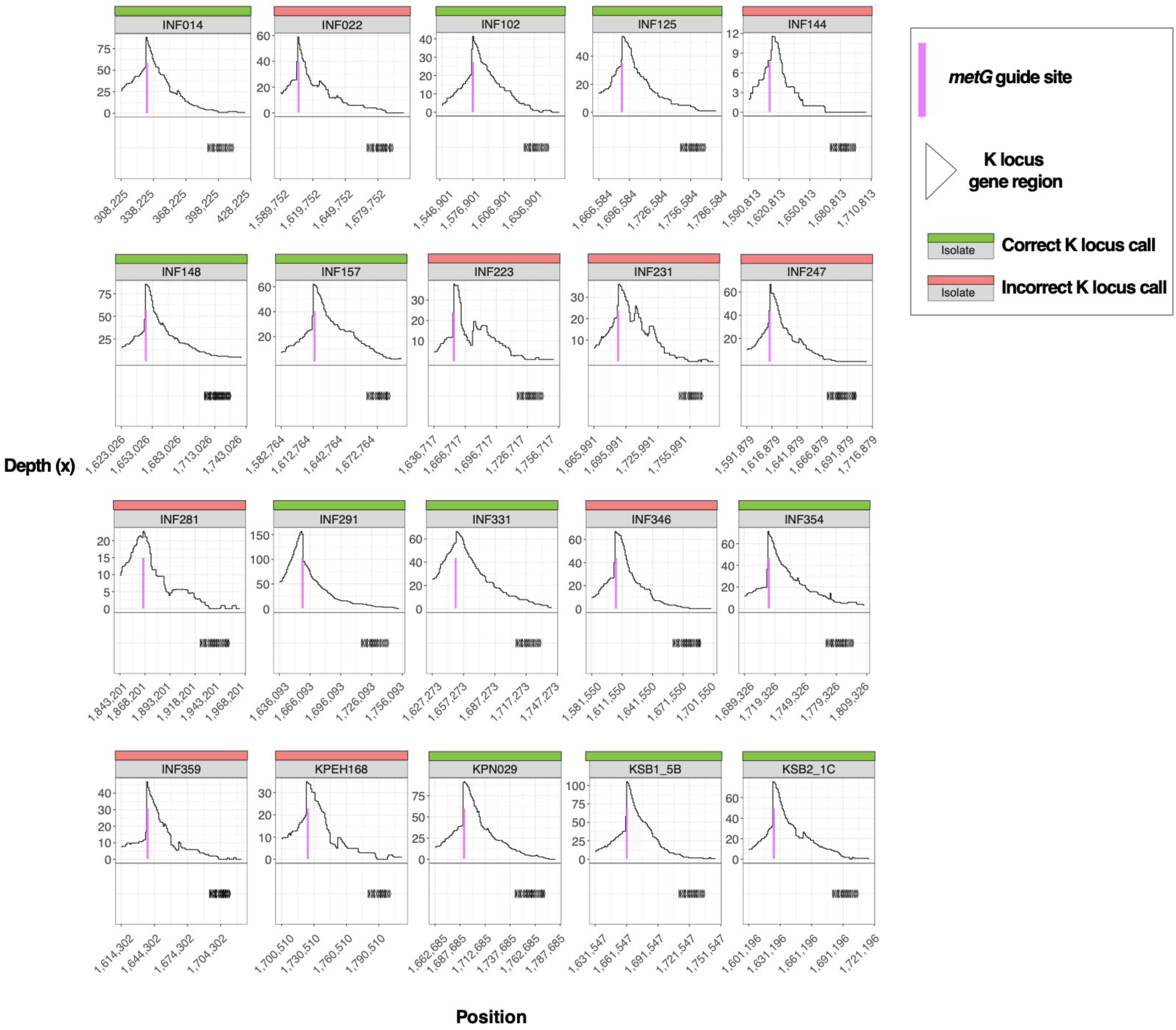


**Supplementary Figure 11 – Depth of ontarget reads aligned against the region surrounding the K locus across *K. pneumoniae* isolates following CRISPR-Cas9 enrichment and ONT sequencing.** Guide target regions are shown in coloured rectangles. On the bottom panel is the regions of K locus genes.

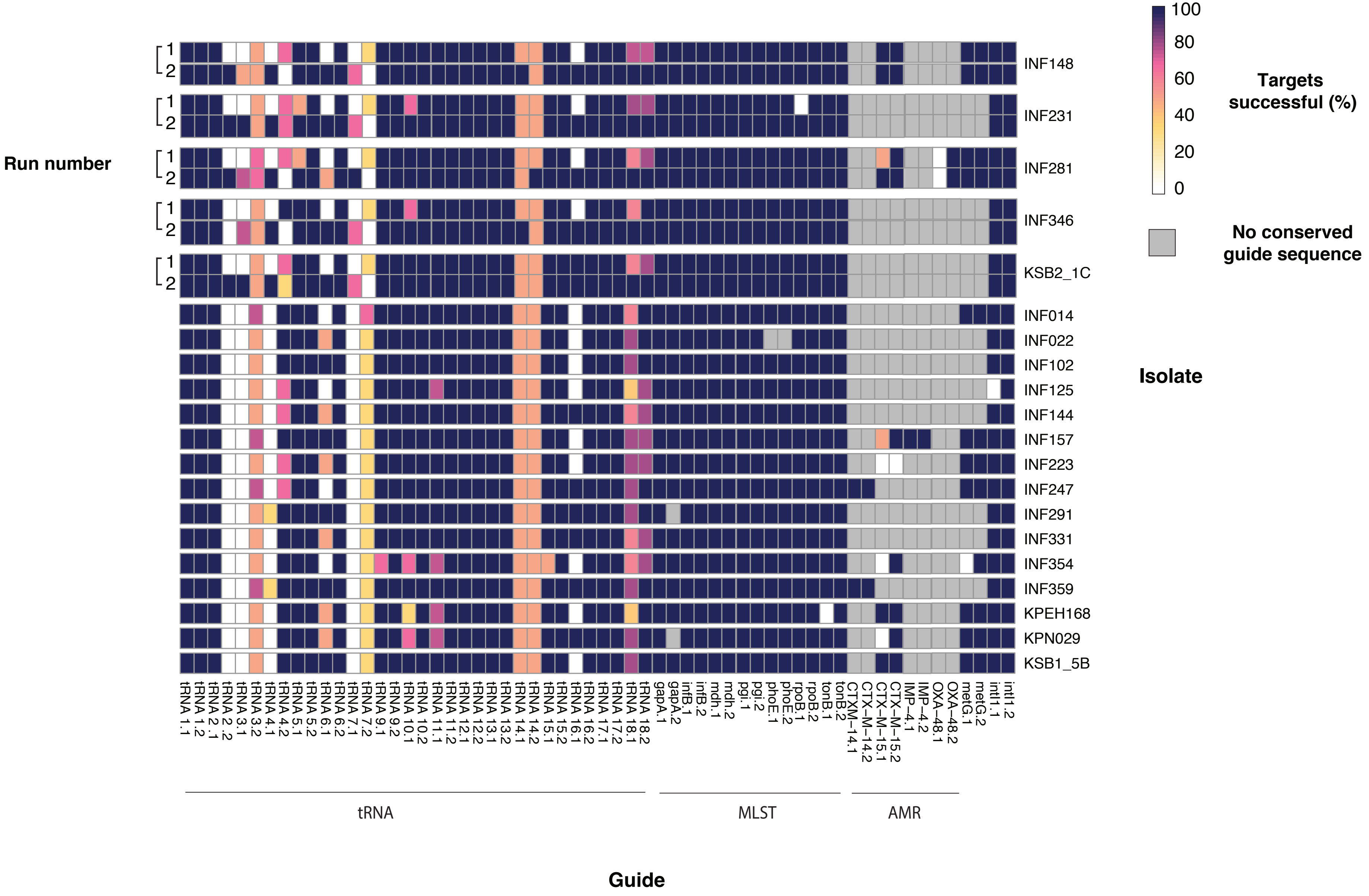


**Supplementary Figure 12 – Individual guide performance across CRISPR-Cas9 enriched libraries of 20 *K. pneumoniae* species complex isolates when using one member of each pair at a time.** A successful target is defined as when the number of ontarget reads is equal to or greater than 10x median depth of offtarget reads. Run number refers to whether it was the initial experiment (run 1) or the repeat experiment (run 2).

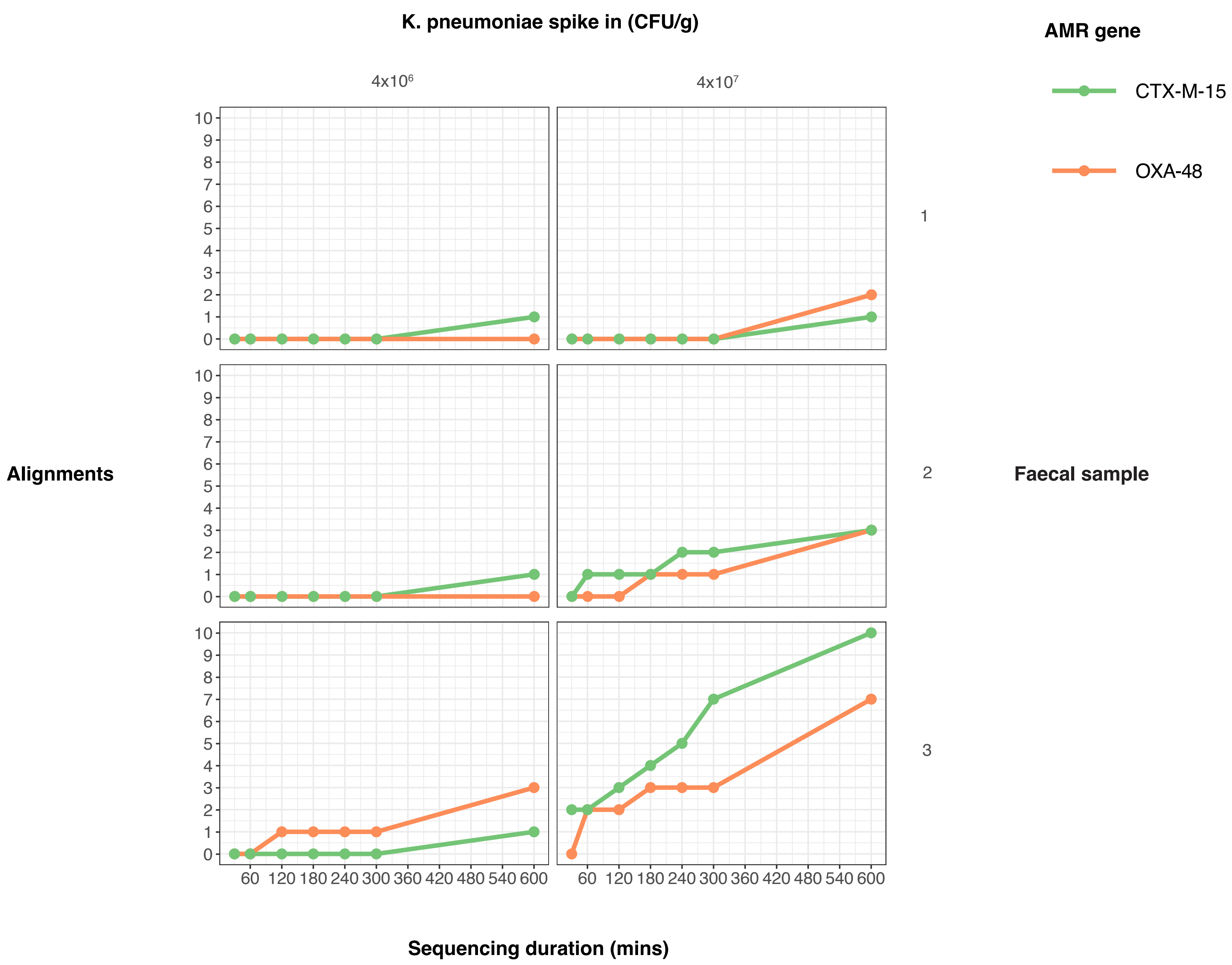


**Supplementary Figure 13 – Alignments to target AMR genes in the first ten hours of sequencing following CRISPR-Cas9 enrichment of three human faecal samples spiked with *K. pneumoniae* at 4 x $10^6$ – $10^7$ CFU/g**

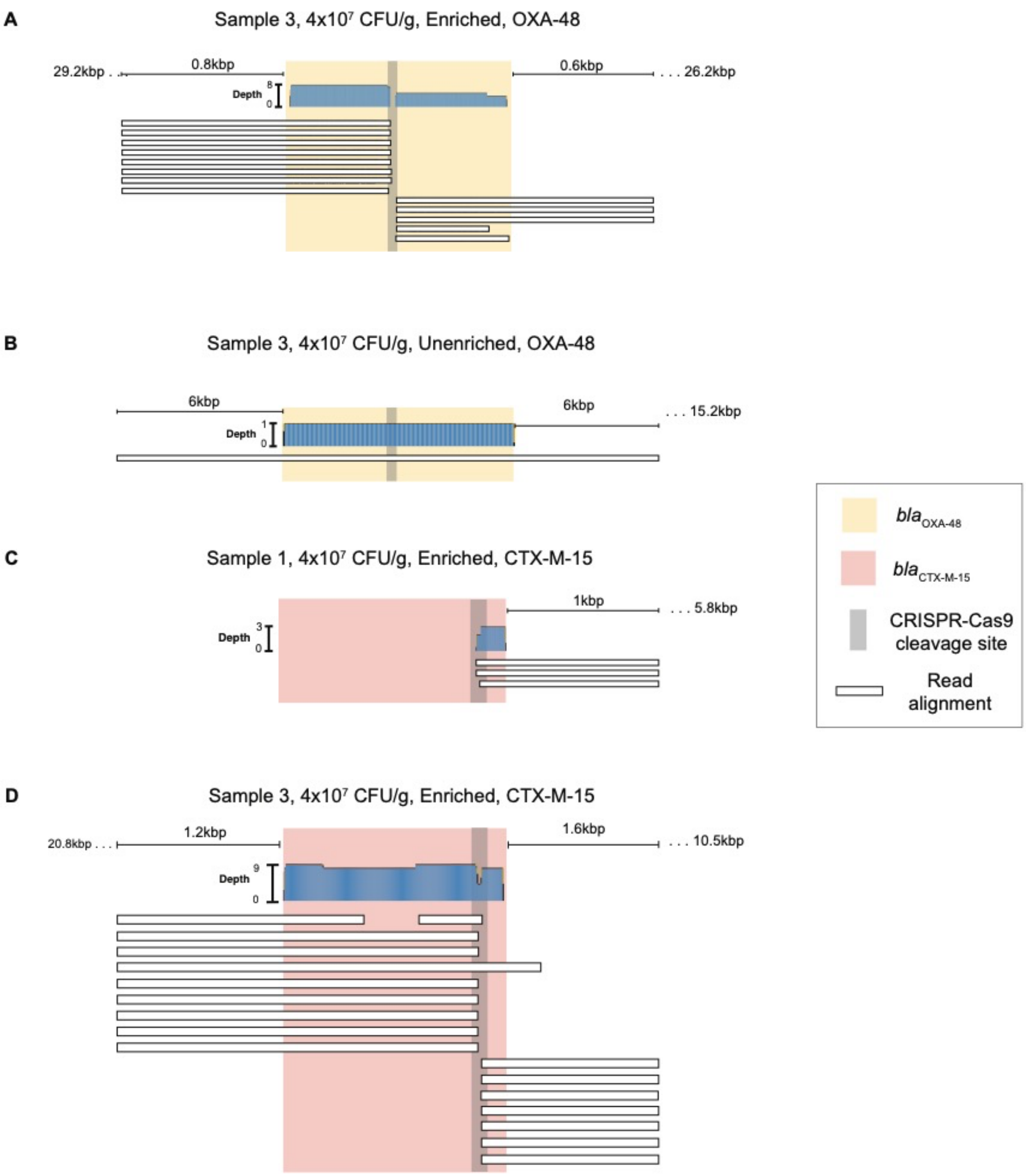


**Supplementary Figure 14 – Read alignment over target AMR genes following sequencing of faecal samples spiked with *K. pneumoniae* strain INF298.**

**A)** Read alignments to *bla*$_{OXA-48}$ after enriched sequencing of sample 3 spiked with INF298 at 4x$10^7$ CFU/g.

**B)** Read alignments to *bla*$_{OXA-48}$ after unenriched sequencing of sample 3 spiked with INF298 at 4x$10^7$ CFU/g.

**C)** Read alignments to *bla*$_{CTX-M-15}$ after enriched sequencing of sample 1 spiked with INF298 at 4x$10^7$ CFU/g.

**D)** Read alignments to *bla*$_{CTX-M-15}$ after enriched sequencing of sample 3 spiked with INF298 at 4x$10^7$ CFU/g.

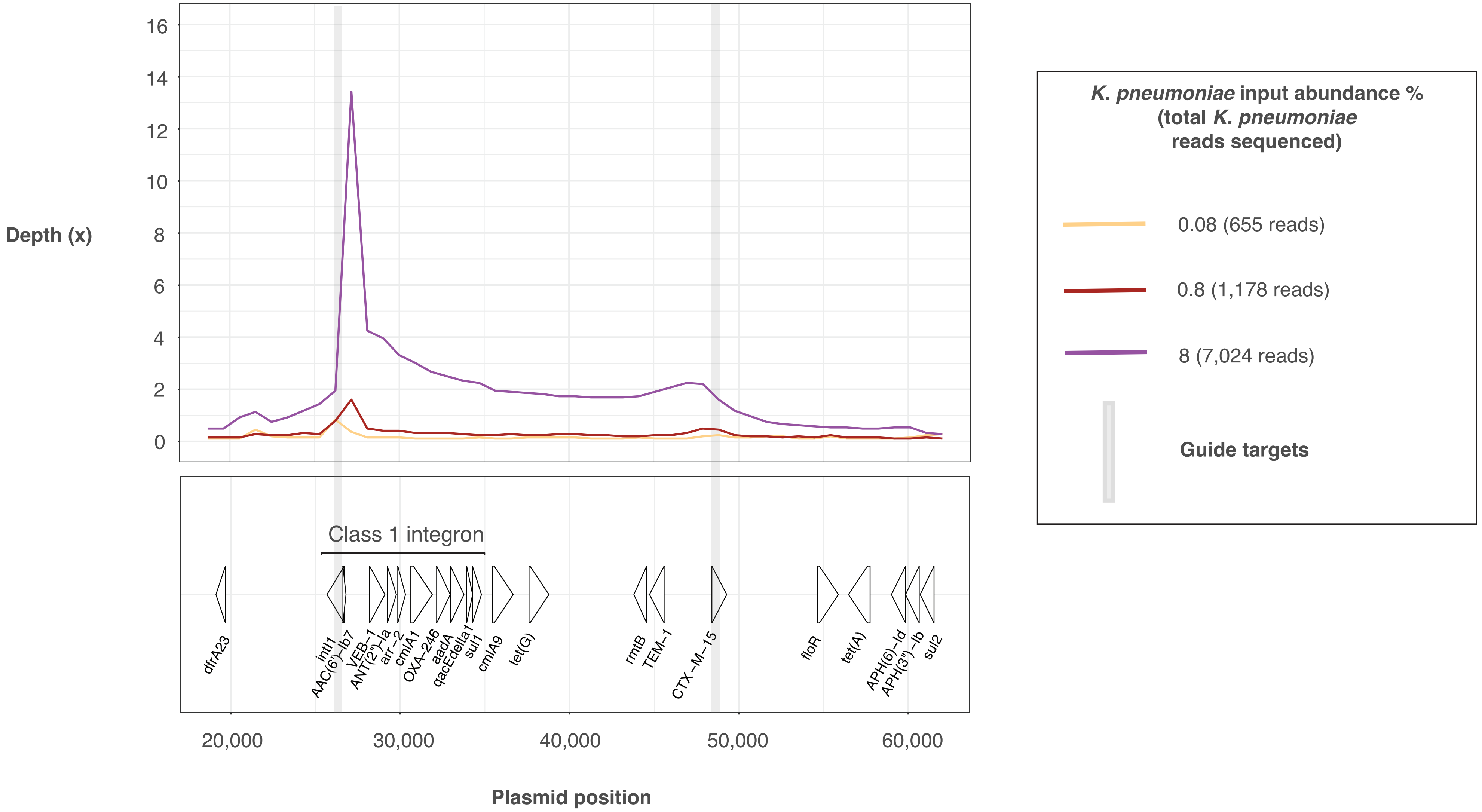


**Supplementary Figure 15 – Depth of an enriched plasmid following sequencing of a mock microbial mixture spiked with *K. pneumoniae* strain INF298 DNA at 0.8 – 8% relative abundance.**

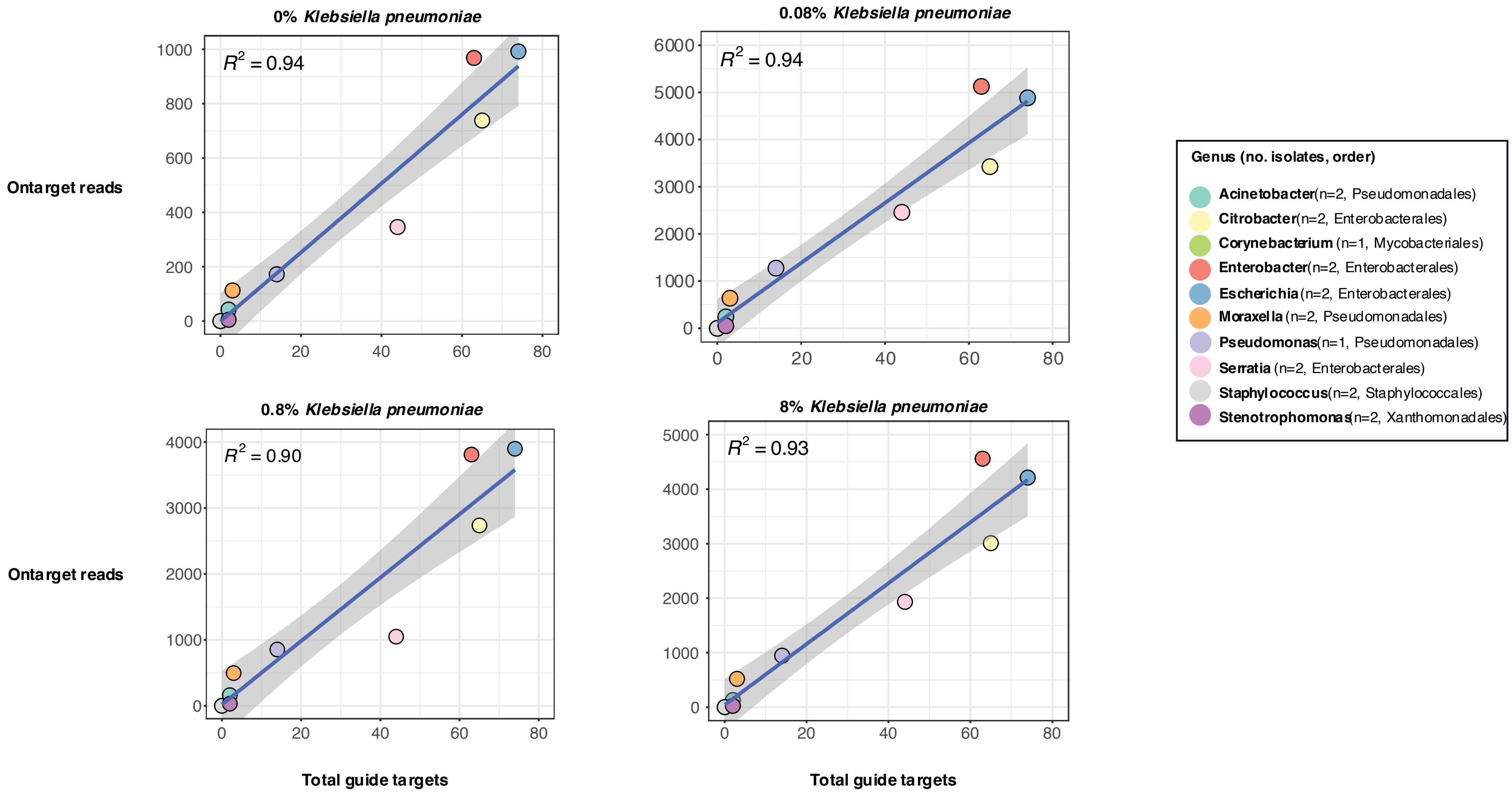


**Supplementary Figure 16 – Relationship between guide conservation and enrichment success amongst isolates in a bacterial community mixture.** Plots show a linear regression between the total guide targets and total ontarget reads for isolates of a given genus. All isolates were included in the mixture at the same abundance except for *K. pneumoniae*, which was included at abundances labelled on top of each plot. *K. pneumoniae* statistics were excluded from linear regression analyses due to different abundance between libraries.

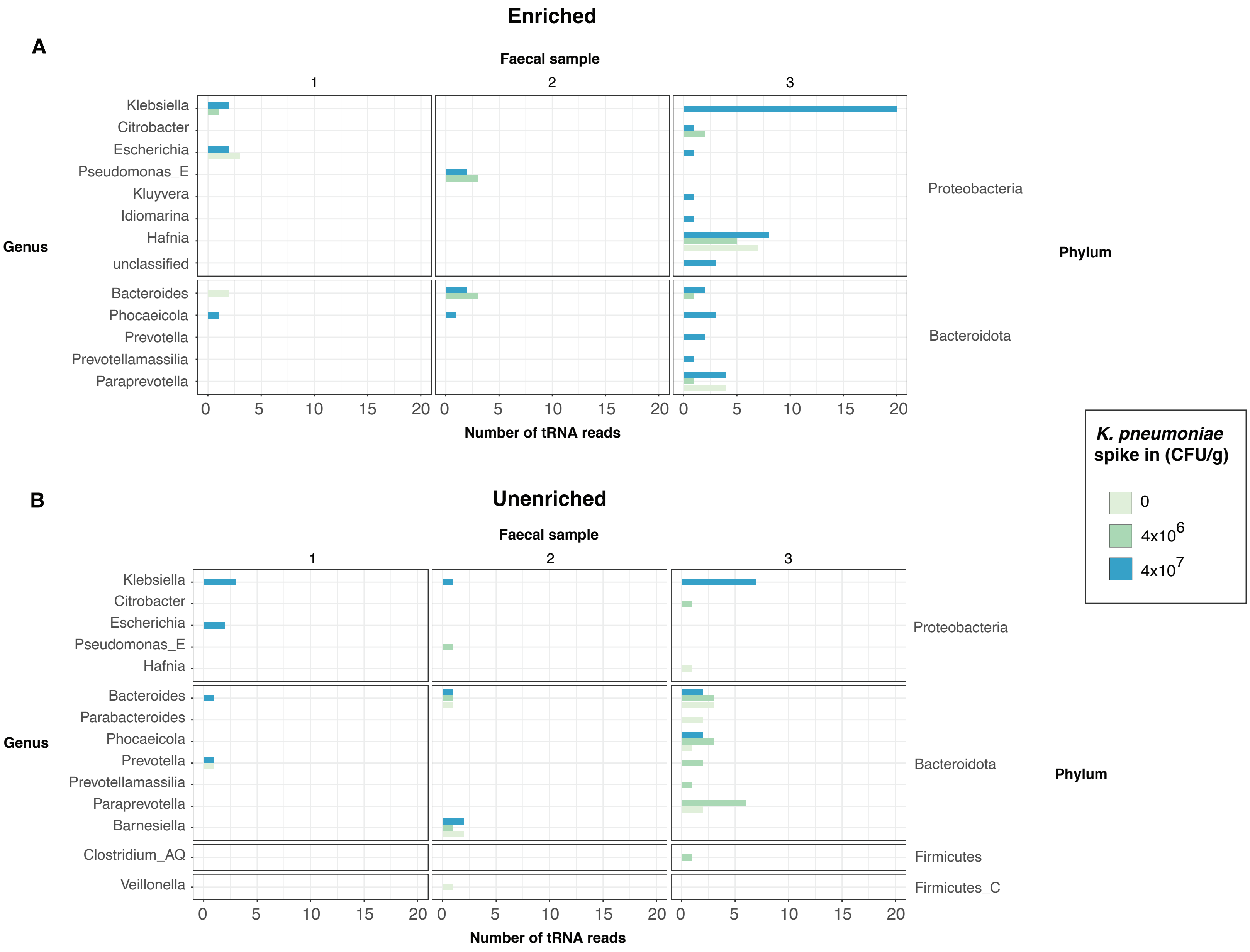


**Supplementary Figure 17 – Taxonomic classification of reads aligning to target tRNA genes following sequencing of three human faecal samples (Parks et al. 2018; Wood et al. 2019).**

**A)** Classification of reads aligning to tRNA genes following CRISPR-Cas9 enriched sequencing. **B)** Classification of reads aligning to tRNA genes following unenriched sequencing**.**
