## Supplemental Methods 1 for "Targeted sequencing of *Enterobacterales* bacteria using CRISPR-Cas9 enrichment and Oxford Nanopore Technologies"

**ONT Sequencing using Cas9 enriched libraries**

**Preparing Cas9 enriched libraries**

**Preparing crRNA guides**

1) crRNA guides come as lypholised pellets, pulse spin before opening tube
2) Prepare 100 uM solution of guide by resuspending in NFTE (pH 7.5), volume for resuspension is oligo yield (nmol on tube label) multiplied by 10
3) Once guides are resuspended keep on ice or at -80oC

**Preparing crRNA guide pool**

1) Mix equal volumes of all guides required for a single pool
2) Mix well by vortexing
3) Aliquot the guide pool into single use tubes (3 ul)
4) Store at -80oC

**Preparing Cas9 ribonucleoprotein complexes (Cas9 RNPs)**

1) Thaw NEB CutSmart Buffer (B7204S) and IDT Duplex buffer (11-01-03-01), vortex and place on ice
2) Combine the following in a PCR tube, this will be sufficient Cas9 RNPs to make 10 libraries if making more than 10 make two lots of Cas9 RNPs
   - 8 ul Duplex buffer
   - 1 ul crRNA pool (100 uM, equimolar)
   - 1 ul tracrRNA (IDT 1072532)
3) Mix by pipetting and spin down
4) In PCR machine heat to 95oC for 5 mins and return to RT for 5 mins
5) Combine to following in 1.5 microfuge tube

- 10 ul of annealed crRNA/tracrRNA (from step above)
- 10 ul NEB CutSmart buffer
- 79.2 ul NFW
- 0.8 ul HiFi Cas9

6) Mix thoroughly by flicking tube
7) Incubate at RT for 30 mins
8) Place on ice

**Dephosphorylating gDNA**

1) Combine the following in a PCR tube
   - 3 ul NEB CutSmart buffer
   - 24 ul of HMW gDNA (at ~210 ng/ul total required ~5 ug) (have used as little as 1 ug of gDNA total and the library still worked)
2) Mix gently by flicking tube
3) Add 3 ul NEB Quick calf intestinal phosphatase (M0525)
4) Mix gently by flicking tube and spin down
5) In PCR machine heat to 37oC for 10 mins, then 80oC for 2 mins and the 20oC hold

**Cleaving and dA tailing gDNA**

1) Thaw dATP (NEB N0440S)
2) Dilute dATP by mixing the following in 1.5 ml microfuge tube
   - 1 ul 100mM dATP
   - 9 ul NFW
3) Vortex to mix and spin down
4) To the PCR tube containing the dephosphorylated gDNA from step above add the following
   - 10 ul Cas9 RNPs
   - 1 ul 10 mM dATP
   - 1 ul NEB Taq polymerase (M0273)
5) Mix gently by flicking tube, spin down
6) In PCR machine heat to 37oC for 60 mins, then 72oC for 5 mins and the 4oC hold (for the guides we have 60 mins is ideal, may need to reduce time is off target cleavage observed)

**Native barcode ligation**

1) For each sample to be barcoded prepare the following mastermix
   - 5 ul NFW
   - 3 ul Native barcode
   - 50 ul Blunt/TA Master Mix
2) Add 30 ul of mastermix to the 42 ul of prepared DNA and mix by flicking tube
3) Add remaining 28 ul of mastermix to the DNA sample and mix by flicking tube
4) Incubate RT 10 minutes
5) Add 50 ul of resuspended AMPure beads and mix by flicking
6) Incubate 10 mins RT
7) Place on magnet, once clear remove supernatant
8) Add 200 ul fresh 70% Ethanol, then remove ethanol, repeat ethanol wash
9) Spin down tube, place back on magnet and pipette of any residual ethanol
10) Remove from rack and add 7 ul NFW
11) Incubate RT 10 minutes
12) Place on magnet
13) QUANTUS 1 ul of individual barcoded samples
14) Pool samples together trying to equalise amounts from samples with variable concentrations (want pooled volume of 65 ul)
15) QUANTUS 1 ul of pooled sample

**Adapter ligation**

1) Thaw T4 Ligase Buffer and Adapter mix (AMII)
2) Tip mix T4 Ligase Buffer
3) Transfer prepared DNA to 1.5 ml microfuge tube
4) Combine the following reagents in 1.5 ml microfuge tube
   - 20 ul T4 Ligase Buffer
   - 10 ul NEBNext Quick T4 DNA Ligase
   - 5 ul AMII
5) Mix by pipetting thoroughly
6) Add 20ul of ligation mix to prepared DNA
7) Mix gently by flicking tube
8) Add remaining 15 ul of ligation mix to prepared DNA/ligation reaction
9) Mix gently by flicking tube, spin tube
10) Incubate reaction 10 mins RT

**AMPure XP bead purification**

1) Thaw tubes of long fragment buffer (LFB) and elution buffer (EB)
2) Add 100 ul TE (pH 8.0) to ligation reaction
3) Vortex AMPure XP beads well to resuspend
4) Add 60 ul AMPure XP beads to Ligation reaction
5) Mix gently by flicking tube, spin tube
6) Incubate reaction 10 mins RT
7) Place on magnet for 5 mins
8) Remove supernatant
9) Take off magnet
11) Add 250 ul LFB and Mix gently by flicking tube
10) Place on magnet for 5 mins
11) Remove supernatant
12) Take off magnet
12) Add 250 ul LFB and Mix gently by flicking tube
13) Remove supernatant
14) Briefly centrifuge
15) Place on magnet
16) Remove all remaining supernatant
17) Take off magnet
18) Add 13 ul EB
19) Mix gently by flicking tube, make sure beads go into solution
20) Incubate RT 10 mins
21) Place on magnet for 5 mins
22) Remove 12 ul of supernatant and place in 1.5 ml microfuge tube
23) Test concentration of library on QUANTUS
    - Expected range 50-150 ng/ul
    - Have run libraries with 5-20 ng/ul, they are not great but still worth running

**Priming and loading flow cell**

1) Thaw SQB, LB, FLT and FB
2) Flick mix all reagents and store on ice until required
3) Add 30 ul of FLT to FB and pipette to mix
4) Open flow cell priming port
5) Remove air bubbles from primimg port by removing 20-30 ul of storage buffer from port
6) Load 800 ul of the combined FB/FLT into priming port, close priming port and wait 5 mins
7) Prepare library by mixing the following in 1.5 ml microfuge tube
   - 25 ul SQB
   - 13 ul LB (mix immediately before pipetting to ensure resuspended)
   - 12 ul library
8) Open priming port and spot on sample port
9) Load 200 ul of FB/FLT into priming port
10) Mix prepared library by gentle pipetting
11) Add 50 ul of library to spot on sample port, dropwise allowing each drop to be adsorbed before adding the next
12) Close spot on sample port and priming port
13) Start run on MinKnow software

**Starting basecalling in Guppy**

1) Choose NO basecalling
2) Open Terminal window
3) CTRL C to kill anything running in Terminal
4) In Finder navigate to the FAST5 parent folder
5) Right click on folder
6) Select open in terminal
7) In terminal type basecalling and then up arrow to retrieve previous commands
8) Once basecalling has started can see;
   - Barcode distribution- opportunity to top up with more pooled or single libraries if necessary
   - Translocation speed- opportunity to refuel it speed drops below 300 bp/s

**Nuclease Flush of a flow cell**

1) When you have collected enough sequence data but there is still life left in the flow cell
2) Stop the run and save the pdf report
3) Thaw a tube of homemade DNAse buffer (380 ul)
4) Add 20 ul of DNAse I to the buffer
5) Open priming port
6) To remove air from system pipette 20-30 ul buffer out of priming port
7) Load 400 ul DNAse solution into priming port
8) Close priming port
9) Incubate RT 30 minutes
10) If using flow cell again immediately proceed with new flush (FLB/FT)
11) Remove all solution from the waste reservoir
12) If storing flow cell add 500 ul Solution S from Wash kit to the priming port
13) Close priming port
14) Remove all solution from the waste reservoir
15) Store flow cell at 4oC
