## Supplemental Methods 2 for "Targeted sequencing of *Enterobacterales* bacteria using CRISPR-Cas9 enrichment and Oxford Nanopore Technologies"

**CRISPR enrichment barcoding**

**Dephosphorylating gDNA**

1) Combine the following in a PCR tube
   - 3 ul NEB CutSmart buffer
   - 24 ul of HMW gDNA (at ~210 ng/ul total required ~5 ug) (have used as little as 1 ug of gDNA total and the library still worked)
2) Mix gently by flicking tube
3) Add 3 ul NEB Quick calf intestinal phosphatase (M0525)
4) Mix gently by flicking tube and spin down
5) In PCR machine heat to 37oC for 10 mins, then 80oC for 2 mins and the 20oC hold

**Cleaving and dA tailing gDNA**

1) Thaw dATP (NEB N0440S)
2) Dilute dATP by mixing the following in 1.5 ml microfuge tube
   - 1 ul 100mM dATP
   - 9 ul NFW
3) Vortex to mix and spin down
4) To the PCR tube containing the dephosphorylated gDNA from step above add the following
   - 10 ul Cas9 RNPs
   - 1 ul 10 mM dATP
   - 1 ul NEB Taq polymerase (M0273)
5) Mix gently by flicking tube, spin down
6) In PCR machine heat to 37oC for 60 mins, then 72oC for 5 mins and the 4oC hold (for the guides we have 60 mins is ideal, may need to reduce time is off target cleavage observed)

**Native barcode ligation**

1) For each sample to be barcoded prepare the following mastermix
   - 5 ul NFW
   - 3 ul Native barcode
   - 50 ul Blunt/TA Master Mix
2) Add 30 ul of mastermix to the 42 ul of prepared DNA and mix by flicking tube
3) Add remaining 28 ul of mastermix to the DNA sample and mix by flicking tube
4) Incubate RT 10 minutes
5) Add 50 ul of resuspended AMPure beads and mix by flicking
6) Incubate 10 mins RT
7) Place on magnet, once clear remove supernatant
8) Add 200 ul fresh 70% Ethanol, then remove ethanol, repeat ethanol wash
9) Spin down tube, place back on magnet and pipette of any residual ethanol
10) Remove from rack and add 14 ul NFW
11) Incubate RT 10 minutes
12) Place on magnet
13) QUANTUS 1 ul of individual barcoded samples
14) Dilute DNA as required, want 13uL for next step

**Regular ONT barcoding**

**End prep**

1) Combine the following reagents in a tube:
   - 1.5-2 ug gDNA in 51 ul NFW
   - 6 ul NEBNEXT Ultra II End-prep reaction buffer
   - 3 ul NEBNEXT Ultra II End-prep enzyme mix
2) Incubate 5 mins RT
3) Vortex the AMPureXP to fully resuspend the beads
4) Add 60 ul AMPureXp beads to each of the samples
5) Tip mix
6) Incubate 5 mins RT
7) Place plate on magnet for 5 mins
8) Remove supernatant
9) Add 200 ul 70% Ethanol to each sample
10) Remove ethanol and repeat wash
11) Spin tube
12) Remove all residual ethanol with a 50 ul pipette
13) Remove tube from magnet
14) Add 11 ul NFW to each sample
15) Tip mix
16) Incubate 10 mins RT
17) Place tube on magnet for 5 mins
18) Measure gDNA concentration on QUANTUS

**Barcode Ligation**

1) Combine the following reagents:
   - 10 ul End-prepped gDNA
   - 2.5 ul Native barcode
   - 12.5 ul Blunt/TA Ligase Master Mix
2) Tip mix
3) Incubate 10 mins RT
4) Vortex the AMPureXP to fully resuspend the beads
5) Add 50 ul AMPureXP beads to each of the samples
6) Tip mix
7) Incubate 5 mins RT
8) Place plate on magnet for 5 mins
9) Remove supernatant
10) Add 200 ul 70% Ethanol to each sample
11) Remove ethanol and repeat wash
12) Spin tube
13) Remove all residual ethanol with a 50 ul pipette
14) Remove plate from magnet
15) Add 14 ul NFW to each sample
16) Tip mix
17) Incubate 10 mins RT
18) Place plate on magnet for 5 mins
19) Measure gDNA concentration on QUANTUS

**Adapter ligation (same for CRISPR and regular)**

**Adapter ligation**

1) Thaw T4 ligase buffer, T4 ligase and Adapter mix (AMII)
2) Tip mix T4 ligase buffer
3) Combine the following reagents in 1.5 ml microfuge tube
   - 13uL DNA
   - 4 ul Ligase buffer
   - 2 ul NEBNext Quick T4 DNA Ligase
   - 1 ul AMII
4) Mix by pipetting thoroughly
5) Add 5ul of ligation mix to prepared DNA
6) Mix gently by flicking tube
7) Add remaining 3 ul of ligation mix to prepared DNA/ligation reaction
8) Mix gently by flicking tube, spin tube
9) Incubate reaction 10 mins RT

**AMPure XP bead purification**

1) Thaw tubes of short fragment buffer (SFB) and elution buffer (EB)
2) Add 20 ul TE (pH 8.0) to ligation reaction
3) Vortex AMPure XP beads well to resuspend
4) Add 12 ul AMPure XP beads to Ligation reaction
5) Mix gently by flicking tube, spin tube
6) Incubate reaction 10 mins RT
7) Place on magnet for 5 mins
8) Remove supernatant
9) Take off magnet
10) Add 75 ul SFB and Mix gently by flicking tube
11) Place on magnet, remove supernatant once clear
12) Repeat SFB wash
10) Briefly centrifuge
11) Place on magnet
12) Remove all remaining supernatant
13) Take off magnet
14) Add 24/num samples +1 uL EB
15) Mix gently by flicking tube, make sure beads go into solution
16) Incubate RT 10 mins
17) Place on magnet for 5 mins
18) Test concentration of library on QUANTUS
    - Expected range 50-150 ng/ul
    - Have run libraries with 5-20 ng/ul, they are not great but still worth running
19) Dilute barcodes in elution buffer separately as required for barcode balancing
20) Pool elutions of each barcode into final volume of 24uL
21) Store in the fridge or proceed to loading the library

**Priming and loading flow cell**

1) Thaw SQB, LB, FLT and FB
2) Flick mix all reagents and store on ice until required
3) Add 30 ul of FLT to FB and pipette to mix
4) Open flow cell priming port
5) Remove air bubbles from primimg port by removing 20-30 ul of storage buffer from port
6) Load 800 ul of the combined FB/FLT into priming port, close priming port and wait 5 mins
7) Prepare library by mixing the following in 1.5 ml microfuge tube
   - 37.5 ul SQB
   - 25.5 ul LB (mix immediately before pipetting to ensure resuspended)
   - 24 ul library
8) Open priming port and spot on sample port
9) Load 200 ul of FB/FLT into priming port
10) Mix prepared library by gentle pipetting
11) Add 50 ul of library to spot on sample port, dropwise allowing each drop to be adsorbed before adding the next
12) Close spot on sample port and priming port
13) Start run on MinKnow software
